# Integrated Approach On Degaradtion Of Azo Dyes Using Laccase Enzyme And Nanoparticle With Its Interaction By *In Silco* Analysis

**DOI:** 10.1101/677690

**Authors:** Rajalakshmi Sridharan, Veena Gayathri Krishnaswamy, Archana Murali.K, Revathy Rajagopal, Thirumal Kumar. D, George Priya Doss.C

## Abstract

Azo dyes, released by the textile industries causes severe damage to the environment and living organisms. The degradation of azo dyes is widely studied using enzymatic methods. Laccase, is a copper containing enzyme that degrades the azo dyes into less toxic compounds. In this work, Laccase enzyme produced by the alkaliphile *Pseudomonas mendocina* in the degradation of mixed azo dye showed 0.386 U/Ml activity at pH 8.5. Combination of enzymatic and green synthesised nanoparticle were used in the degradation mixed azo dye. Laccase used in the degradation of mixed azo dyes showed 58.46% in 72 hours while the photocatalytic degradation of mixed azo dyes showed 15.98%. The degradation of azo dyes using copper iodide nanoparticle resulted in 15.835% of mixed azo dye degradation. But it was noticed that combined method removed 62.35% of mixed azo dyes in 60 minutes. Interaction of laccase enzyme with azo dyes using in silico analysis predicted the binding energy of RR (−7.19 kcal/mol), RB (−8.57 kcal/mol) and RBL (−9.17 kcal/mol).

## INTRODUCTION

The increase in urbanization and population leads to increase in use of dyes, plastics and chemicals that add major pollutants to the environment. Dyes are complex structural compounds that are used in textile, food and pharmaceutical industries (Shyamala Gowri *et al.*, 2014). Azo dyes are synthetic dyes that are used to give permanent colour to fabrics as they are highly resistant to external factors. More than 70% of textile industries uses azo dyes for dying fabrics. Its effluent released into the environment poses a major contribution to environmental pollution (Saratale *et al.*, 2011). The azo dye contains chromophore azo group (N=N) which gives colour to the fabrics. The azo dyes may be classified as mono azo, diazo or poly azo dye based on number of azo groups present in it (Harshad *et al.*, 2015). These azo dyes are released into the environment either untreated or partially treated. This leads to pollution of soil, water and other ecosystems as azo dyes are carcinogenic and mutagenic in nature (Kumar Raven and Sumangala Bhat, 2012). The phytoplankton, zooplankton, fishes, aquatic plants are affected by decreased oxygen level, decreased photosynthetic activity which results in death of aquatic habitats. It also affects human beings by causing jaundice, tumour, skin irritation, allergies, heart defects etc… (Montaya *et al.*, 2015, Salleh *et al.*, 2011, Vakili *et al.*, 2014). These effects have drawn interest in treating the azo dye contaminated sites by various physical, chemical and biological methods. Physical methods such as adsorption and chemical methods such as Fenton‘s reagent and other techniques such as Advanced Oxidation Process (AOPs), Cavitation, Ultrasound technique were studied exclusively (Harshad *et al.*, 2015). The physical and chemical methods are highly toxic and hence biological methods is an alternative eco-friendly technique for treatment of azo dyes (Kobayashi *et al.*, 1982, Saratale *et al.*, 2011).

The biological process involves aerobic and anaerobic degradation of azo dyes by microorganisms. The extremophiles mainly alkaliphiles has drawn interest to study the degradation of azo dyes. As the nature of azo dye contaminated sites are alkaline, it makes the survival of mesophiles difficult. Alkaliphiles are microorganisms whose cell wall is composed of acid polymers which maintains the internal pH of the microorganisms and makes them survive in extreme alkaline environments (Sujatha *et al.*, 2017, Alain *et al.*, 2002). The degradation of azo dyes by microbes occurs through enzymatic cleavage. The enzymatic degradation of azo dyes by the bacteria occurs either by extracellular or intracellular enzymes. The enzymes that are commonly involved in azo dye degradation are Azoreductase, Laccase, Peroxidases etc… These enzymes require appropriate environmental conditions to maintain its stability and activity (Joshni *et al.*, 2011). Laccase are copper containing oxidases that does not require redox mediators for degrading azo dye (Zahra *et al.*, 2014 and Luciana Pereira *et al.*, 2009). The substrate specificity of laccase can be widened by addition of redox mediators to the reaction mixture (Fabbrini *et al.*, 2002). Laccases cleave azo dyes and forms nitrogen molecules that are non-toxic (Andrea *et al.*, 2005).

The photocatalytic degradation is one among the Advanced Oxidation Process (AOPs) which is used in bulk treatment of pollutants. This process results in the mineralization of the recalcitrant compounds (Comparelli *et al.*, 2005). Recently, nano-size materials have been widely studied because of their better properties compared to the bulk size particle. They exhibit a large specific surface area and thus have distinct, catalytic and thermal properties. Copper iodide is a highly versatile compound having many applications in adsorption studies, catalysis, solar cells, etc (Tavakoli *et al.*, 2013). The green synthesis of copper iodide nanoparticles gives highly pure product involving the use of minimum chemicals. Photo-assisted catalysis using a semiconductor has been recognized as a promising approach for the elimination of many organic pollutants, such as azo dyes (Anushya Vijayakumar, Revathy Rajagopal, 2016).

In this present study, the degradation of individual and mixed textile azo dyes was studied using a Laccase enzyme producing alkaliphilic bacterium isolated from an azo dye contaminated site. Although there are few reports on photo-degradation using green synthesized metal oxides nanoparticles, there is hardly any data on dye degradation involving a combination of green synthesized nanoparticles and bacterial laccase. So, it was thought worthwhile to enhance mixed azo dye degradation by immobilizing bacterial laccase on CuI nanoparticle. The analysis of degradation of azo dyes were done using UV – Visible Spectrophotometer, HPLC, and GC-MS. The dyes were also analysed using *in silico* analysis which showed the binding of laccase enzyme with Reactive Black (RBL), Reactive Red (RR) and Reactive Brown (RB). The binding also correlates with the degradation of the azo dyes.

## MATERIALS AND METHODS

### SAMPLE COLLECTION

The textile dye contaminated soil samples were collected from Erode (textile industry). Three different types of soil (different colour) were collected and used for isolation of azo dye degrading bacterial strains. The textile dyes Reactive Red (RR), Reactive Brown (RB), and Reactive Black (RBL) were purchased from Bengal ChemiColor and Co, Parrys, Chennai, India. All the other chemicals were purchased from Sigma Aldrich, India.

### ISOLATION OF ALKALIPHILIC BACTERIAL STRAIN

The bacterial strains were isolated from the textile effluent contaminated soil samples which were enriched in the Mineral Salts Medium (MSM) with 100 ppm of mixed azo dye. The MSM was prepared using Disodium hydrogen phosphate (Na_2_HPO_4_ – 12.8 g/L), Potassium dihydrogen phosphate (KH_2_PO_4_ – 3 g/L), Ammonium chloride (NH_4_Cl – 1 g/L), Sodium chloride (NaCl – 0.5 g/L), Magnesium sulphate (MgSO_4_ – 10 mg/L), 0.01 M Calcium chloride (CaCl_2_) and 20% glucose (pH 8.5). The medium was sterilized, cooled and and mixed azo dye containing RR, RB, and RBL each of 100 ppm was added in 250 mL Erlenmeyer flask. The bacterial strains were isolated on MSM Agar and the pure culture of each strain was stored in 15 % glycerol at - 20°C for further studies (Tarun and Rachana, 2012).

### DEGRADATION ASSAY

The maximum absorbance of the dye (RR, RB, RBL, and Mixed) was determined using UV Vis spectrophotometer. The culture was provided with 100 ppm of mixed azo dye in MSM and the decolorization was monitored every 24 hours for 4 days. The degradation percentage was calculated using the formula (Andrea *et al.*, 2005).

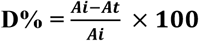

Where D – degradation %, Ai – initial absorbance, At – Absorbance after time t.

### BIODEGRADATION OF AZO DYE DEGRADATION

The biological method for mixed and individual azo dye degradation was carried out using laccase enzyme produced by isolated bacterial strain DY2. The enzymatic degradation was studied further by extraction of extracellular and intracellular proteins. The efficiency of the degradation of mixed and individual azo dyes was studied.

### EXTRACTION OF LACCASE ENZYME

The presence of Laccase enzyme was screened in extracellular and intra cellular extracts of the isolated bacterial strain. The laccase production by the bacterial strain DY2 was determined by inoculating the bacterium in 100mL MSM broth amended with 100 ppm of mixed azo dye. The culture was incubated at 37°C and centrifuged at 10,000 g for 10 min, then the supernatant was filtered and used for further studies (Satish Kalme*et al.*, 2009).The intracellular laccase enzyme was extracted by suspending the pellet in 50 mmol^−1^ of potassium phosphate buffer of pH (7.4), then sonicated using ultra probe sonicator at 60 amp for 4 mins with 1 min interval at 4°C. The supernatant was filtered using 0.44µ Millipore filter. The filtrate was then assayed for laccase enzyme activity (Dawkar *et al.*, 2008).

Laccase activity was determined by 2,2’ -azino-bis (3-ethylbenzothiazoline-6-sulphonic acid) ABTS substrate oxidation. The Volume of 0.5 mL of ABTS (0.1M citrate buffer, pH – 4.5), 0.5 mL of enzyme sample was incubated at 37°C, 120 rpm for 10 min (Zahra *et al.*, 2014). Absorbance was observed at 379 nm by a UV/Vis spectrophotometer (Hitachi, double beam). One unit of enzyme activity was defined as the amount of enzyme required to reduce the absorbance of 0.01/sec (Arka Mukhopadhyay *et al.*, 2013). The laccase enzyme activity in U/mL was calculated using the formula given below

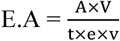

where, E.A – enzyme activity, A-absorbance at 379 nm, V-total volume of a mixture (mL), t-incubation time, e – extinction co-efficient, v – volume of enzyme (mL). The molar extinction co-efficient was calculated and found to be 1.398 M^−1^ cm^−1^ (Alkesh*et al.*, 2017).

### EFFECT OF pH AND TEMPERATURE ON LACCASE

The pH and temperature were optimised to determine the maximum enzyme activity of laccase produced by DY2KVG bacterial strain by varying the pH of buffer and the incubation temperature used in the assay. The laccase activity for pH of various range (5.8 to 8.0) using 50 mM Potassium phosphate buffer and pH of 9.0 using 50 mM Glycine -NaOH buffer were used for optimisation (Mohamed Neifar *et al.*, 2016). The temperature ranging from 20°C to 50°C were used to study the effect of temperature on the laccase activity. The enzyme activity of laccase was monitored using a standard laccase assay procedure after 10 min of incubation (Alkesh *et al.*, 2017).

### AZO DYE DEGRADATION BY PHOTOCATALYSIS

The photocatalytic degradation of azo dyes (RR, RB, RBL, and Mixed dyes) was carried out in a batch reactor of 200 mL capacity. The reactor was fed with 50 mL of MSM at neutral pH and 50 mL of azo dye (RR, RB, RBL, and Mixed dye). The degradation was carried out with UV irradiation and continuous stirring. The UV lamp was placed parallel to the quartz tube. The temperature was maintained at 21°C ± 0.2°C. The samples were withdrawn every 10 mins up to 1 hour and the samples were scanned from 200 – 800 nm (Maulin Shah, 2013).

### EXTRACT PREPARATION FROM *Hibiscus rosa – sinensis* L. FLOWER

The *Hibiscus rosa – sinensis* L. flower was collected from our college premises and washed using distilled water. The petals of the flower were separated and ground to form a paste using distilled water. The paste was further diluted by addition of distilled water and centrifuged, from which the supernatant was used for the synthesis of Copper Iodide nanoparticle (CuI) (Archana *et al.*, 2019). The process was carried out without further purification using AnalR-Grade CuSO4.5H2O and KI and the preparation of CuI nanoparticle form the flower extract was performed based on the literature (Indubala *et al.*, 2018).

### CuI CHARACTERIZATION

The synthesised CuI nanoparticle was characterized using X-ray powder diffraction (Bruker D8 advance P–XRD), scanning electron microscope (Carl Zeiss MA15/EVO18), energy-dispersive analysis of X-ray (Oxford INCA Energy 250 Microanalysis System), Fourier transform infrared spectroscopy (Bruker FTIR spectrometer, Alpha-T), UV–visible spectroscopy (Jasco UV–Vis Spectrophotometer, V-50) (Archana *et al.*, 2019).

### DEGRADATION OF AZO DYE BY COPPER IODIDE (CuI) NANOPARTICLE

The photocatalytic degradation of azo dyes (RR, RB, RBL, and Mixed dyes) was carried out in same reactor used for photocatalysis degradation of azo dyes. In this process of degradation, 25 mg of CuI nanoparticle were added in addition to MSM (50 mL) and azo dye (50 mL) as in photocatalytic degradation (Maulin Shah, 2013).

### COMBINED METHOD FOR DEGRADATION OF AZO DYES

The efficiency of Laccase enzyme to degrade azo dye was increased by immobilizing the Laccase enzyme onto CuI nanoparticle. CuI NP – 10 mg, 1mL of laccase enzyme with 0.341 U/mL of enzyme activity and 1 mL of potassium phosphate buffer (pH - 8) were added and incubated in shaker for 24 hrs. The mixture was centrifuged at 10,000g for 10 mins and washed twice with phosphate buffer of pH 8 (Sanjay *et al.*, 2014). The immobilized laccase enzyme was introduced into the reactor containing an equal volume of azo dyes, MSM and the degradation percentage was calculated (Maulin Shah, 2013).

### ANALYSIS OF METABOLITES AFTER AZO DYE DEGRADATION

The HPLC was carried out using Shimadzu Prominence binary gradient HPLC system (Christ Lab, Stella Maris College), C_18_ column by isocratic using methanol gradient. The samples were prepared using ethyl acetate (equal volume of sample and solvent). The solvent phase was extracted and filtered using membrane filter (0.44µ) and then condensed in rotary vaccum evaporator. Then 20µl of the sample was injected manually into the column with methanol as mobile phase (Harshad *et al.*, 2015). Gas Chromatography-Mass Spectrum was carried out for the sample prepared for HPLC and the metabolites produced by the degradation were identified.

### MOLECULAR IDENTIFICATION OF ISOLATED BACTERIUM

The genomic DNA was isolated from each of the bacterial strains and PCR amplification was performed. The unknown bacterial strains were given for 16S r-RNA sequencing in Amnion Biosequences, Bangalore. The evolutionary relationship between the bacterial strains was determined by constructing a phylogenetic tree using the Maximum Likelihood Method in MEGA7.

#### *In silico* Analysis

The laccase protein sequence of *Pseudomonas mendocina* was retrieved from NCBI database with GenBank ID KER98563.1. This sequence was used to model the 3D structure using Swiss-Model online server with PDB ID 1RW0 as a template (Biasini *et al.*, 2014). The model was evaluated using the Rampage server (Lovell *et al.*, 2003). The active site of the protein was predicted using CastP server (Binkowski, Naghibzadeh, & Liang, 2003). The 3D structures of RR, RBl, RB were retrieved from PubChem database with CIDs 6012481, 5360531, and 166118. Molecular dockings were performed twice using AutoDock standalone package (Morris *et al.*, 2009). The hydrogen and necessary charges were introduced to the laccase protein, and torsions were set to the reactive dyes. The grid of 60×60×60 cubic along x, y, and z-axiswere fixed around the laccase active site. AutoGrid and AutoDock were performed consecutively using Lamarckian Genetic Algorithm. The average binding energy and Laccase-Dyes complexes were obtained as the output. These complexes were visualized using Schrodinger Maestro.

## RESULTS AND DISCUSSION

The azo dyes are degraded using conventional methods of which biodegradation is considered as an eco-friendly method for degradation of azo dyes. The enzymes produced by the bacterial strains degrades the azo dyes based on the nature of the enzyme and the bacterial strain producing it. The present study focuses on the degradation of azo dyes (RR, RB, RBL and mixed azo dye) by laccase enzyme produced by an alkaliphilic bacterial strain. The degradation of azo dyes by photocatalysis and by nanoparticles were also studied. This gave rise to the idea of immobilizing the nanoparticle with the laccase enzyme in azo dyes degradation.

### BIODEGRADATION OF AZO DYES

The azo degrading bacterial strain was enriched using 100 ppm of mixed azo dye a sole carbon source at alkaline pH (8.5) in MSM agar and the isolated bacterium was designated as DY2KVG (Figure 1). The growth of the isolated bacterial strain DY2KVG was monitored for 4 days at 540 nm. The bacterium enters the log phase at 48^th^ hour and the decrease in the growth was observed at 96^th^ hour (Figure 2). The absorbance of the mixed azo dye decreased at 48^th^ hour of degradation by the isolated DY2KVG bacterial strain (Figure 3).The enzyme activity of the extracellular laccase enzyme was found to be 0.1 U/mL at 72 hours which was higher compared to the enzyme activity of intracellular enzyme (0.072 U/mL) produced by the bacterium DY2KVG as in figure 4.Thus, the further studies were carried out using the extracellular laccase enzyme.

**FIGURE 1.**
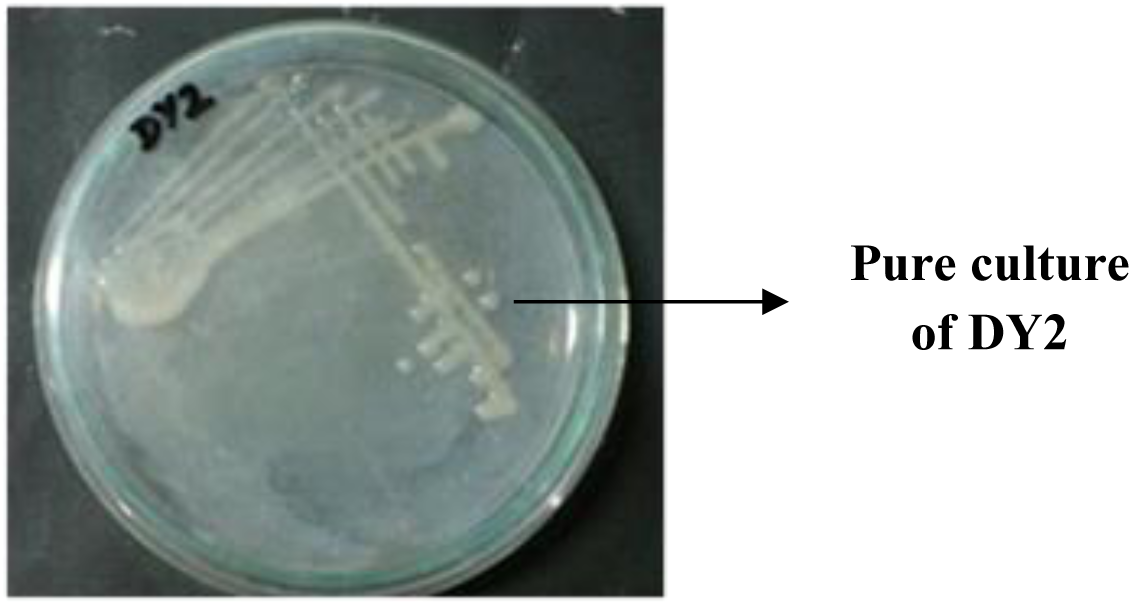
PURE CULTURE - DY2KVG.

**FIGURE 2.**
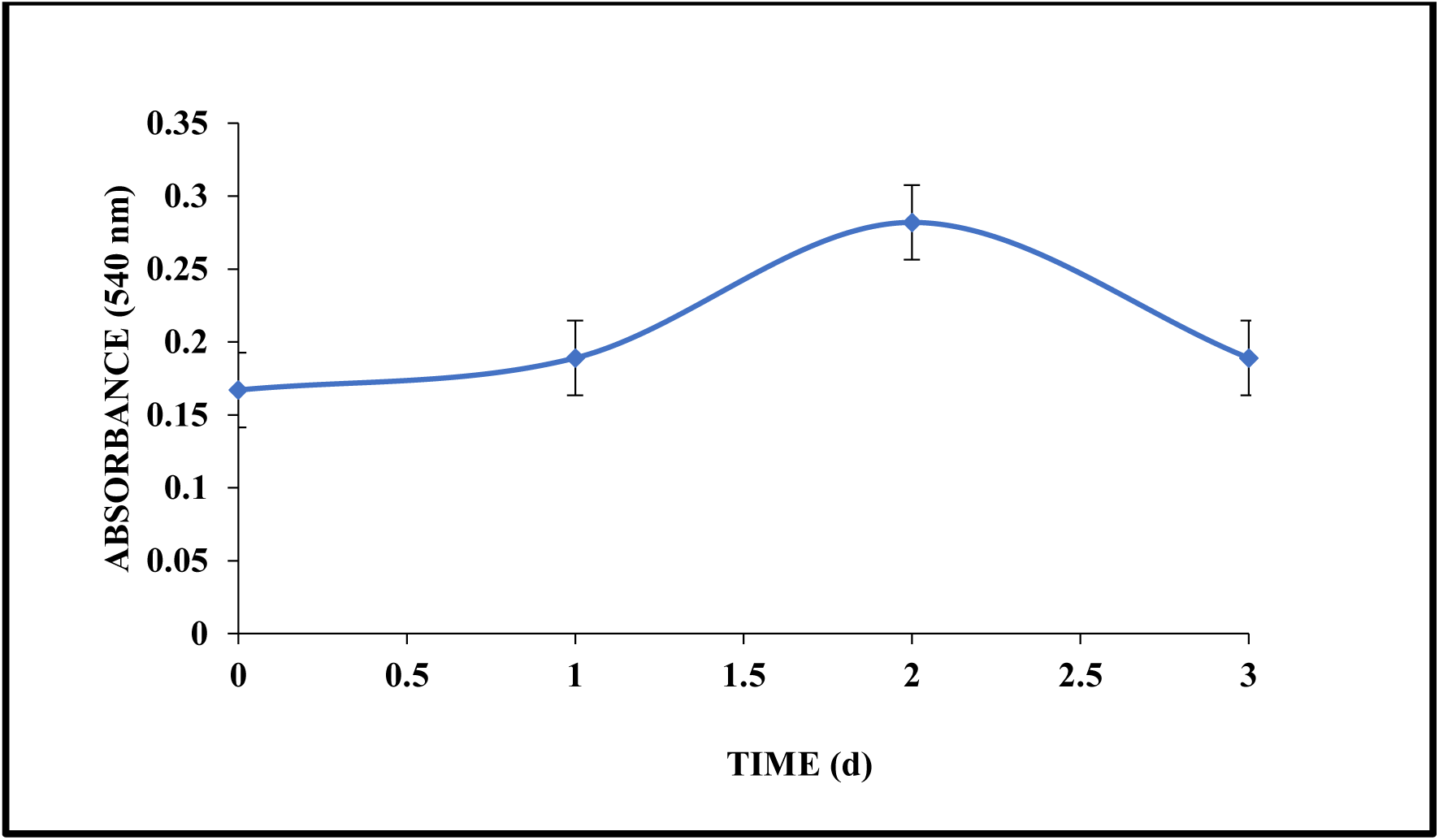
GROWTH CURVE - DY2KVG.

**FIGURE 3.**
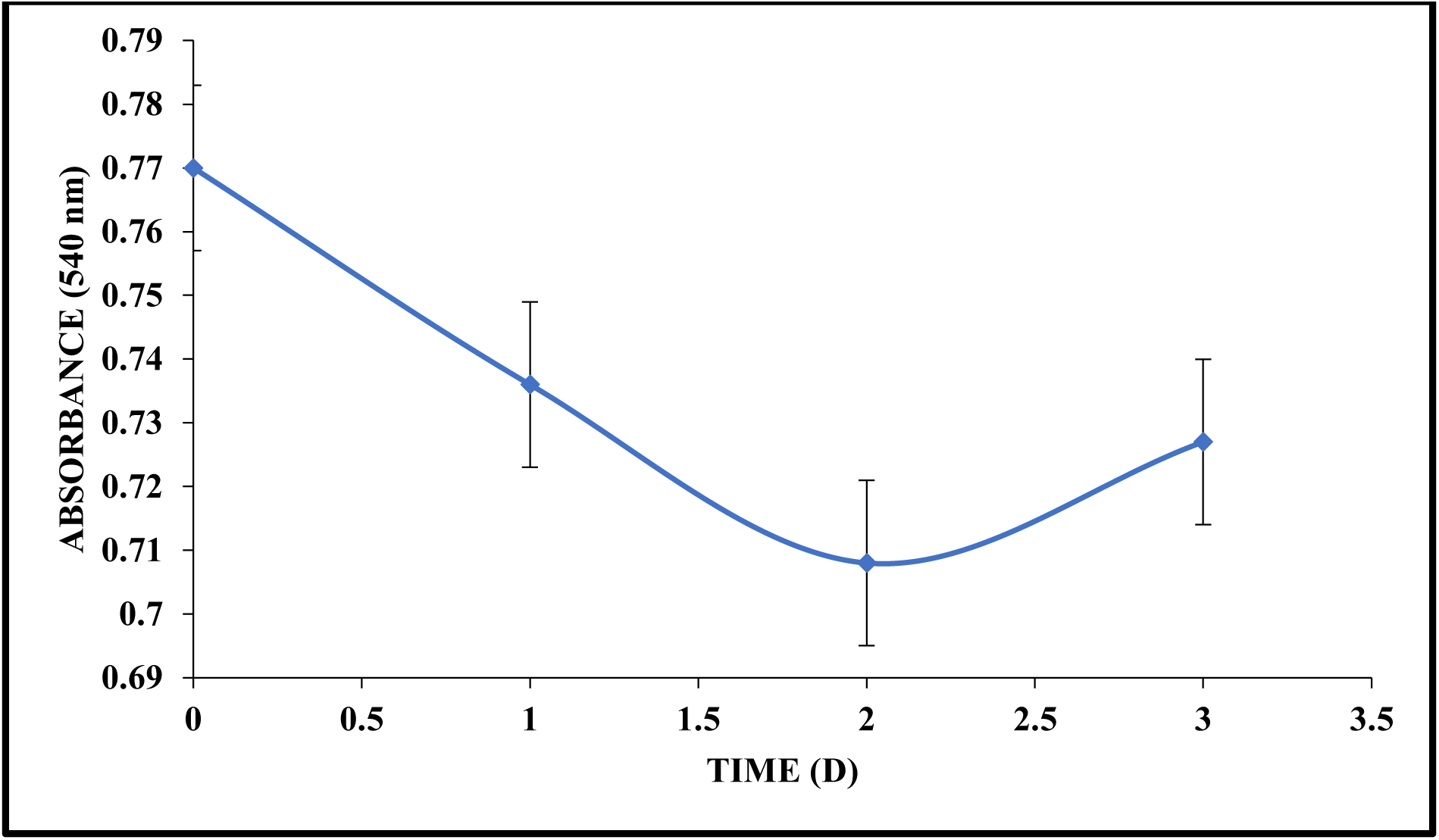
DEGRADATION OF MIXED AZO DYE BY DYE2KVG.

**FIGURE 4.**
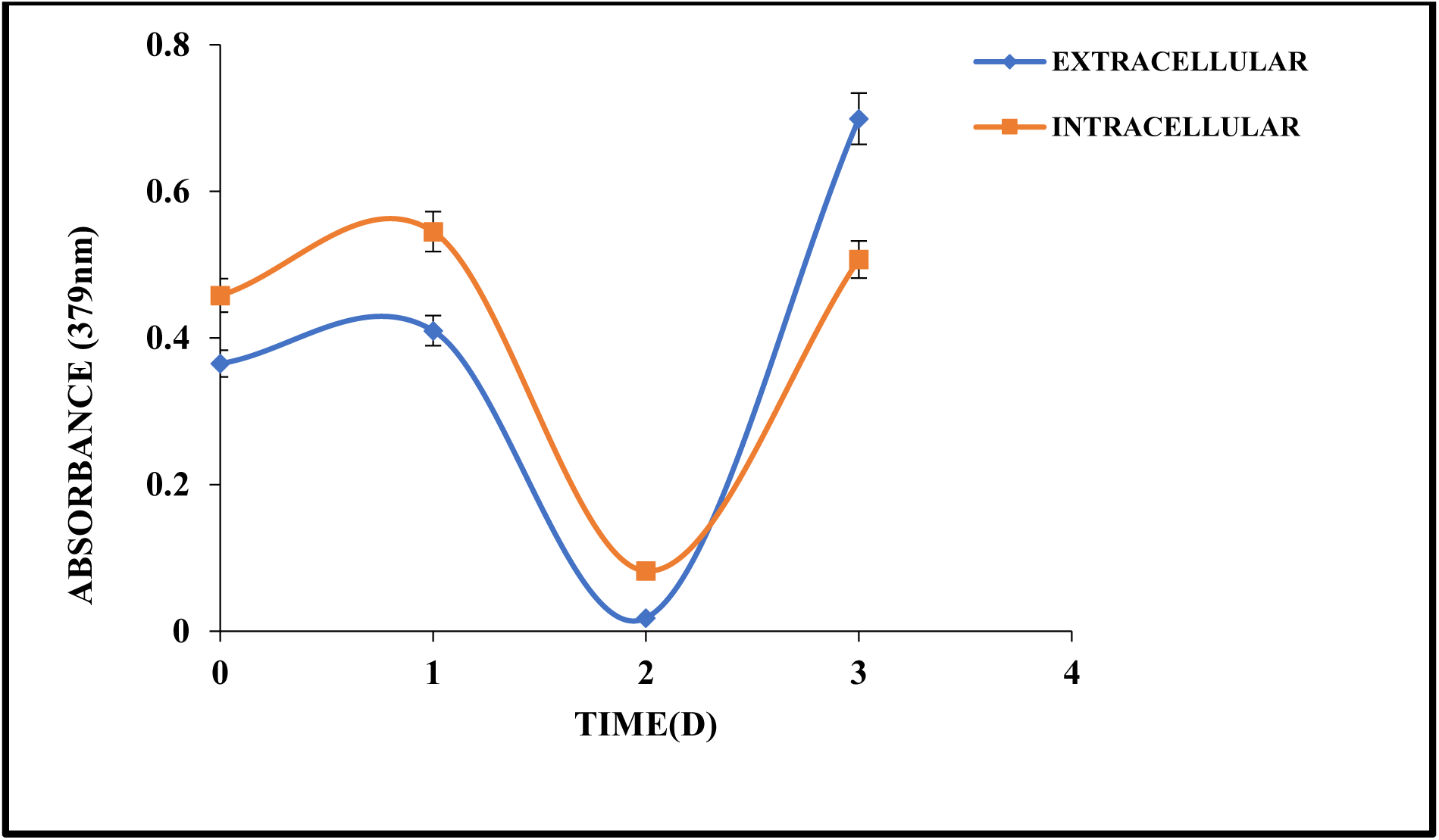
ENZYME ASSAY - LACCASE.

### OPTIMISATION OF pH AND TEMPERATURE

The pH and temperature of the laccase enzyme produced by the bacterial strain isolated from the textile effluent contaminated soil was optimised. The results obtained shows that the laccase enzyme activity increased at pH 5.8 (0.386 U/mL) and decreased with increase in pH. The laccase enzyme activity was found to increase at the pH of 8.0 whose enzyme activity was (0.341 U/mL). This might be due to alkaliphilic nature of the bacterium and the pH of the medium in which the bacterium was isolated (Figure 5). The enzyme activity of laccase enzyme was also monitored at various temperatures. The enzyme activity (0.769 U/mL) was found to be maximum at 20°C and decreased with increase in temperature (Figure 6). The enzyme activity also increased at 50°C (0.6151 U/mL) and the reason for the increase is yet to be identified. Alkesh *et al.*, (2017) isolated laccase enzyme producing *Bacillus sp*. with 0. 981 U/mL of enzyme activity at pH 9 and by addition of peptone as substrate. The extraction of intracellular laccase from *P.desmolyticum* but the optimum pH and temperature for the degradation of Reactive Red and Reactive Green was 4.5 and 60°C (Kalme *et al.*, 2009). The laccase enzyme produced by the *Podoscypha elegans* which resulted in maximum activity in the pH ranging between 5.5 to 7.0 (Satadru Pramanik and Sujata Chaudhuri, 2018) which correlates with the results of present study.

**FIGURE 5.**
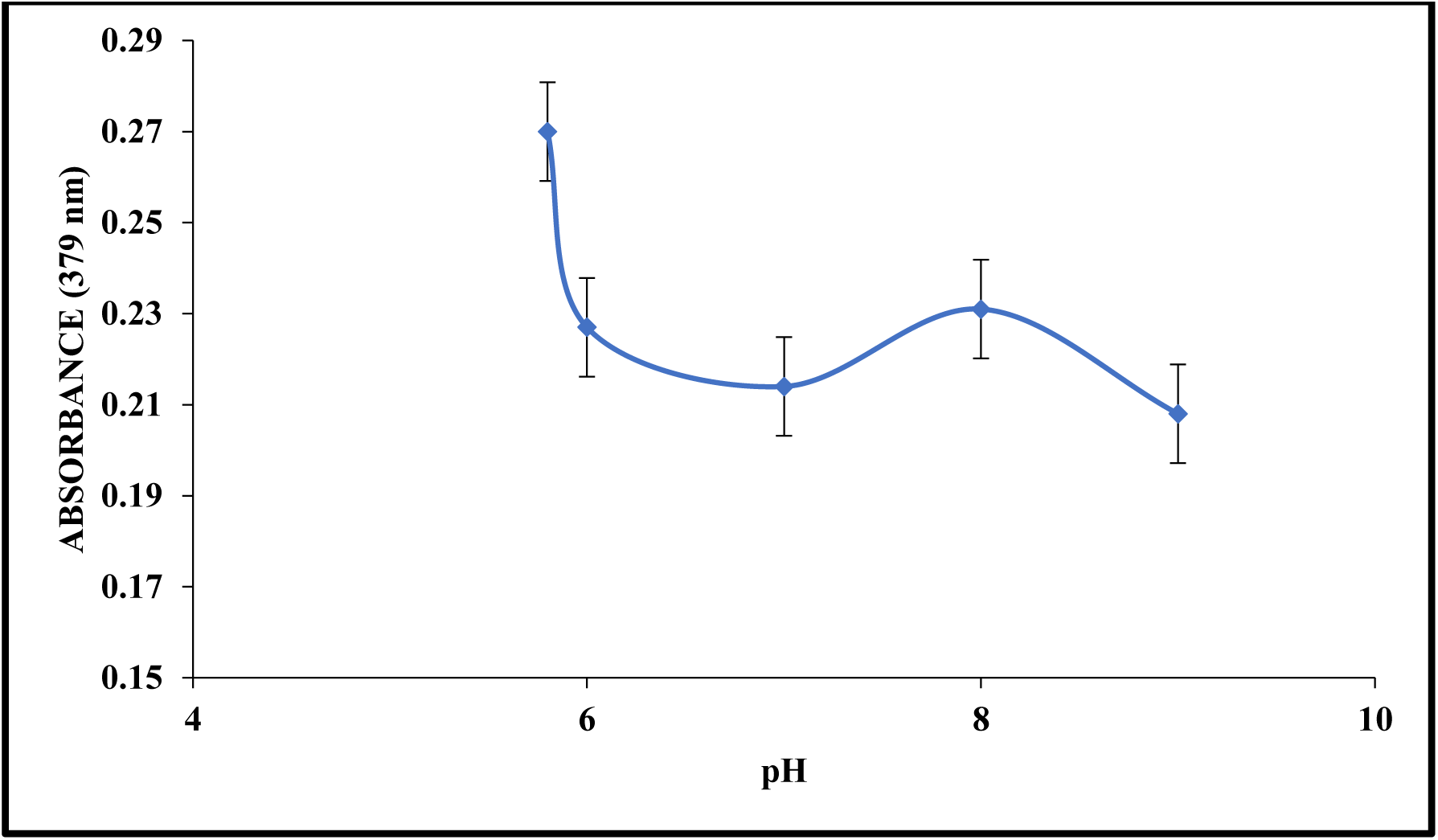
EFFECT OF pH ON LACCASE ENZYME.

**FIGURE 6.**
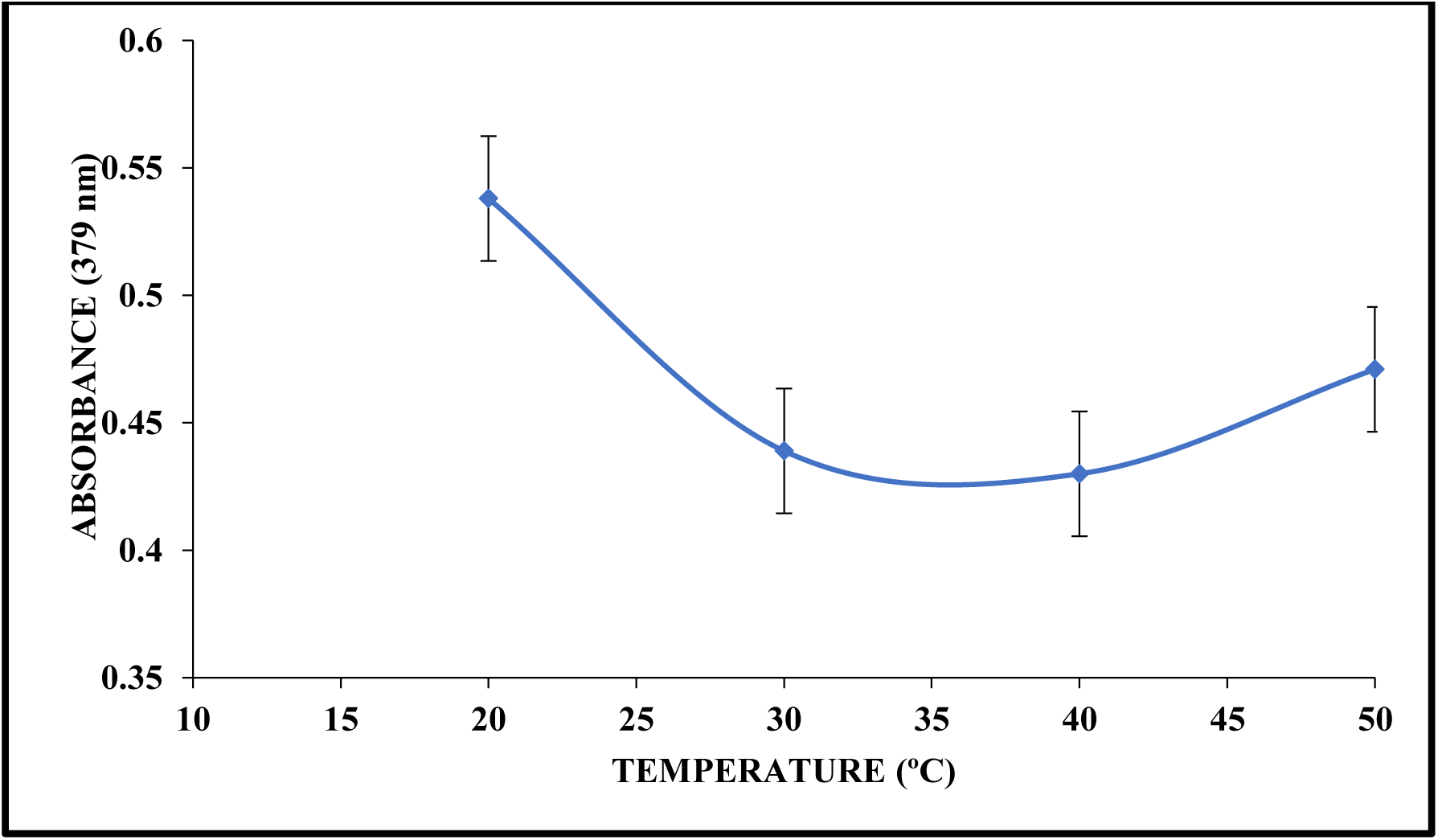
EFFECT OF TEMPERATURE ON LACCASE.

### BIODEGRADATION ANALYSIS OF AZO DYES

In the present study, laccase enzyme produced by an alkaliphilic bacterial strain degraded 58.46 % for mixed dyes, 2.81% for RR, 6.25 % for RBL and 16.47% for RB at the end of 72 hours in single cycle. There was no degradation observed in Direct Blue 15 by the laccase enzyme produced by the fungus *Podoscypha elegans*. The maximum degradation of 70% of Congo red and Rose Bengal was observed in 72 days where as Direct Blue 15 resulted only in 24.8% degradation (Satadru Pramanik and Sujata Chaudhuri, 2018). This contradicts the present study, as the laccase enzyme produced by an alkaliphilic bacterial strain degraded the reactive azo dyes in 72 hours, while the laccase produced by the fungi *Podoscypha elegans* requires 21 days which resulted in minimum degradation of dyes. The bacterial consortium of *P. rettgeri* strain HSL1 and *Pseudomonas sp.* SUK1 showed only up to 22% of 100 mg/L of azo dyes degradation in 48 hours (Harshad *et al.*, 2014).

In this study, the metabolites produced by the degradation of azo dyes by laccase enzyme was analysed by several instrumentation. The HPLC chromatogram of the Mixed Dye (Figure 7), RR (Figure 8), RB (Figure 9) and RBL (Figure 10) dyes showed the decrease in the peak intensity and shift in the retention time of the peaks. Figure 11 shows the GC-MS chromatogram of the metabolites formed by enzymatic degradation of mixed azo dyes after 72 hours. The formation of Aniline, Benzoic acid, Naphthalene, 3 – hydroxyl benzoic acid cyclohexyl 2-buten -1-ol, Catechin and Bendazac methyl ester shows that the complex structure of the parent compounds is broken into many ring structures. Harshad *et al.*, 2015, degraded the Congo red by the bacterial consortium and determined the metabolites by GC-MS. The metabolites were found to be Naphthalene, at RT of 20.55 and Biphenyl compounds after degradation of Congo red. Rakesh *et al.*,2015 study of the degradation of Reactive Red 135 by *P.aeruginosa* were determined by GC-MS, reported naphthalene at RT of 20 and additional compounds such as 2-amino-1-naphthol at 25 RT. Madhuri *et al.*, (2014) reported the degradation products of the direct red dye was Naphthalene and 1,4 – benzediamide. Adosinda *et al.*, (2003) studied the degradation of mixed and individual reactive azo dyes and the by-products were found to be meta-hydroxy benzoic acid at 20.45 retention time.

**FIGURE 7.**
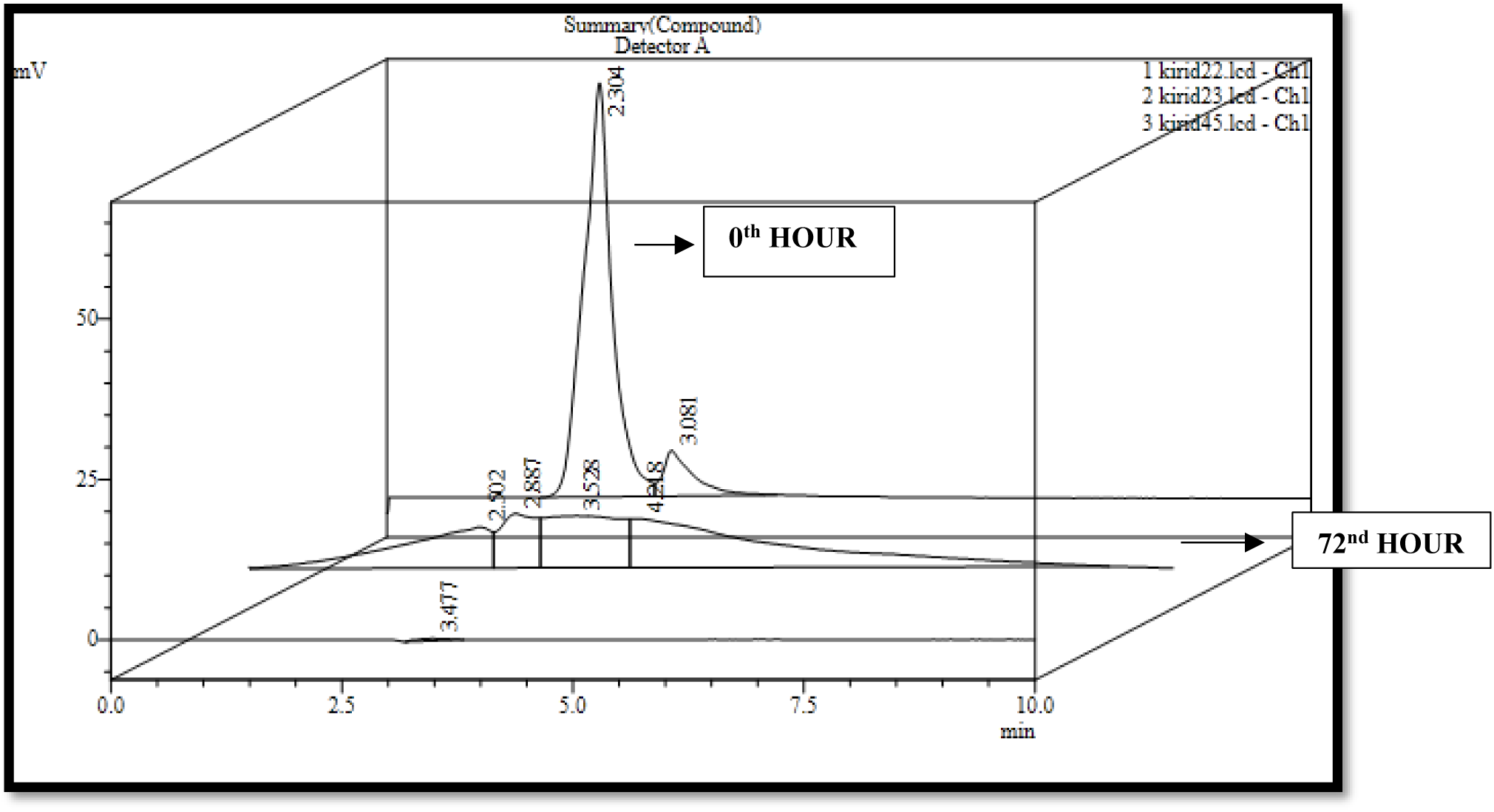
HPLC CHROMATOGRAM OF MIXED AZO DYE.

**FIGURE 8.**
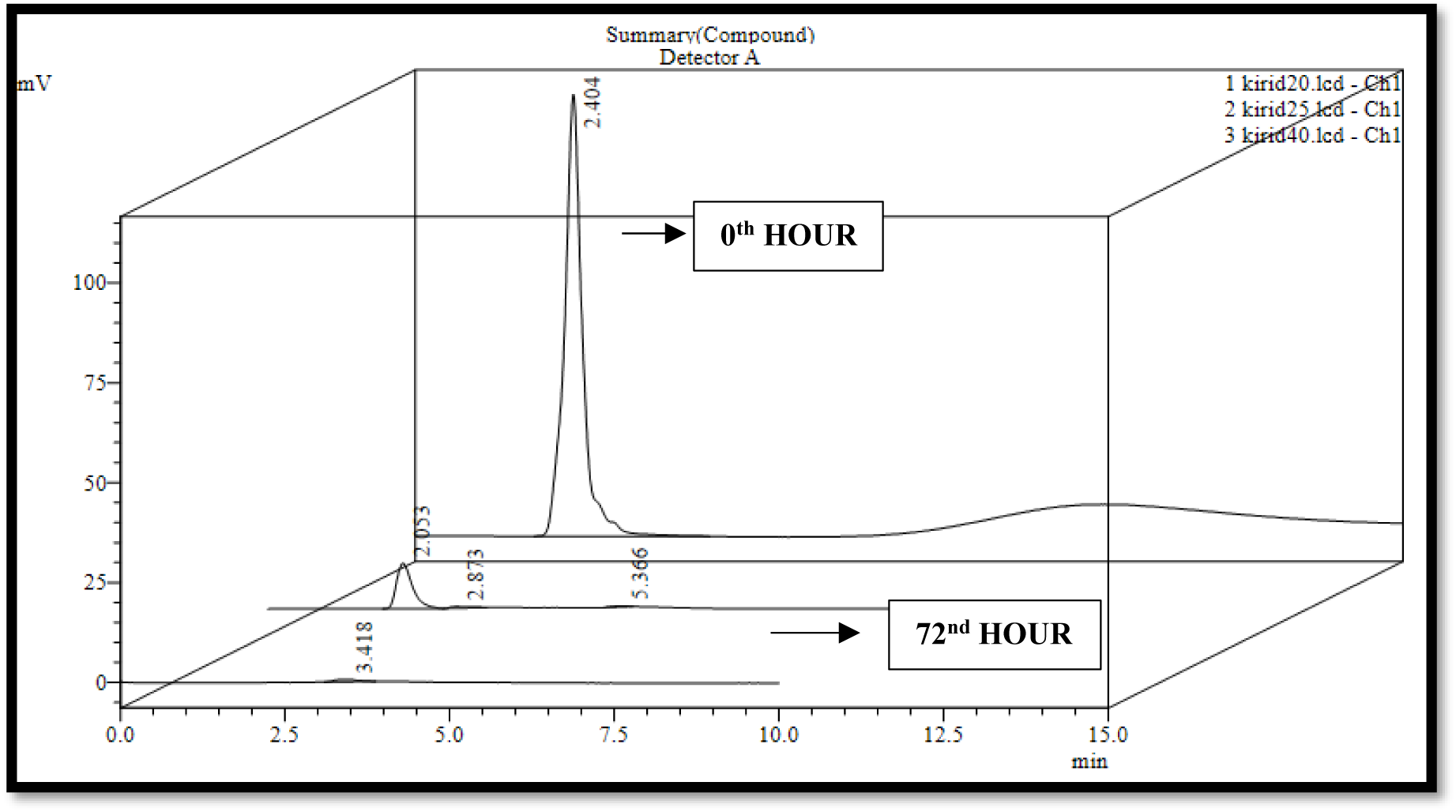
HPLC CHROMATOGRAM OF RR AZO DYE.

**FIGURE 9.**
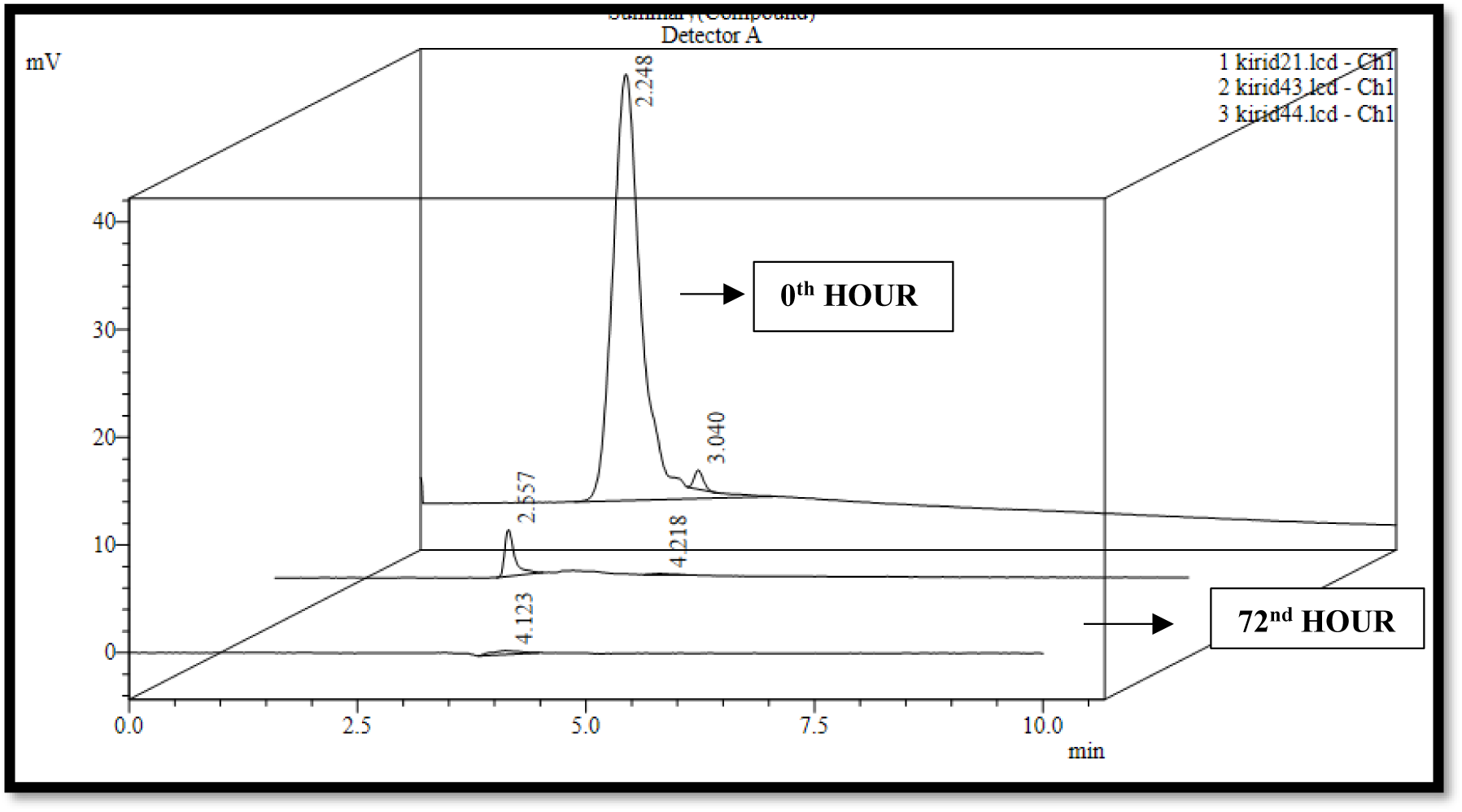
HPLC CHROMATOGRAM OF RB AZO DYE.

**FIGURE 10.**
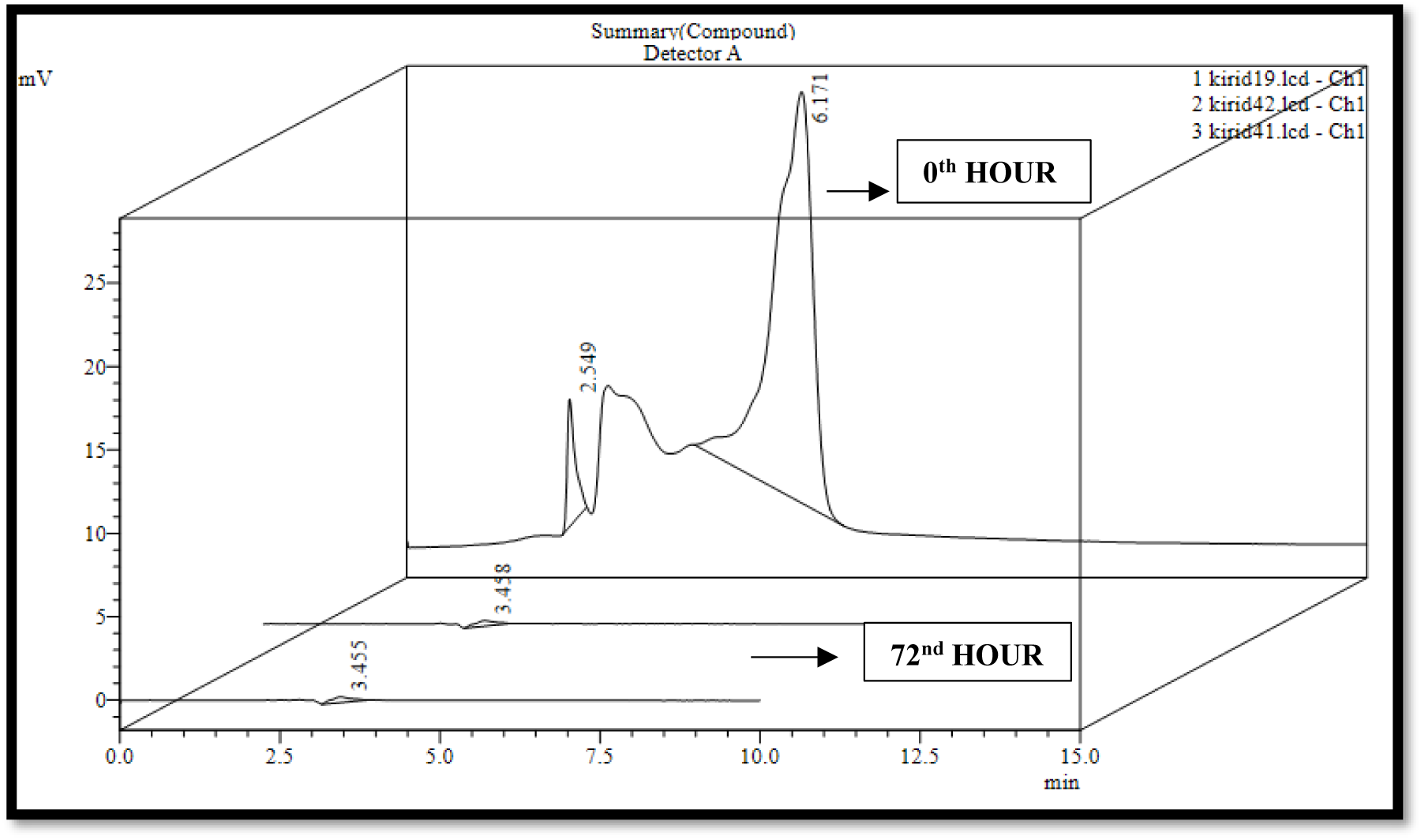
HPLC CHROMATOGRAM OF RBL AZO DYE.

**FIGURE 11.**
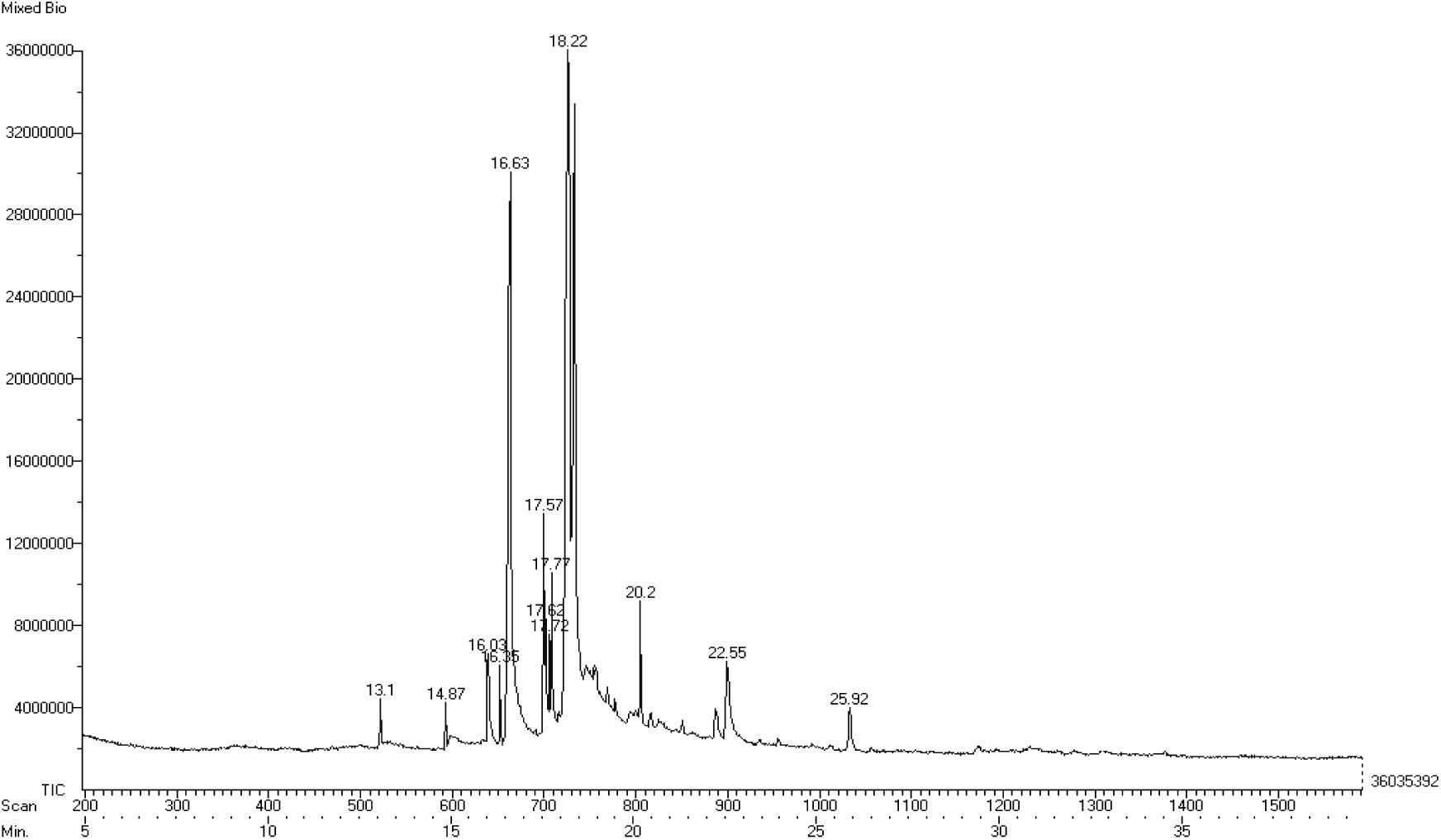
GC-MS - ENZYMATICALLY DEGRADED MIXED AZO DYE.

### PHOTOCATALYSIS OF AZO DYES

The degradation of zo dyes by the photocatalytic breakdown using UV light. This study was performed to determine the effect of UV light in the degradation of azo dyes. The percentage of degradation of azo dyes by the UV light in a photoreactor was observed to be 3.33% for RBL, 15.98% for Mixed dye, 2.694% for RR dye and 23.79% for RB at the end of 1 hour. Thus, the degradation of reactive azo dyes using photocatalytic process was not effective. The process of photodegradation was reported as an effective method for degradation of various organic compounds (Daneshvar *et al.*, 2004). The complete degradation of 50 ppm of Orange II by photocatalysis was reported in 90 min while the increase in concentration increased the time of degradation of orange II (Divya *et al.*, 2013).

### CHARACTERISATION OF CuI NANOPARTICLE

The Copper iodide particles synthesized by green route were characterized by X-ray diffraction (XRD) to determine the structure of the particles (Figure 12). The peak positions (2θ) of the synthesized CuI are 25.64, 29.66, 42.34, 50.07, 52.46, 61.33, 67.51, 69.51 and the XRD pattern was found to match well with that reported in literature (Tavakoli*et al.*, 2013) (JCPDS card no, 82-2111). The diffraction peaks corresponded to (111), (200), (220), (311), (400), (331) and (420) planes of crystalline γ-CuI and was found to fit well in *fcc* system. It was observed that the peaks were sharp, well-defined and no significant impurities were observed in the XRD pattern indicating high purity of the product. The Figure 13 shows morphology and size of the copper iodide particles, which were determined using SEM analysis. In the SEM image, the particle appears to be in triangular shape and the average particle size was found to be 660.92nm.

**FIGURE 12.**
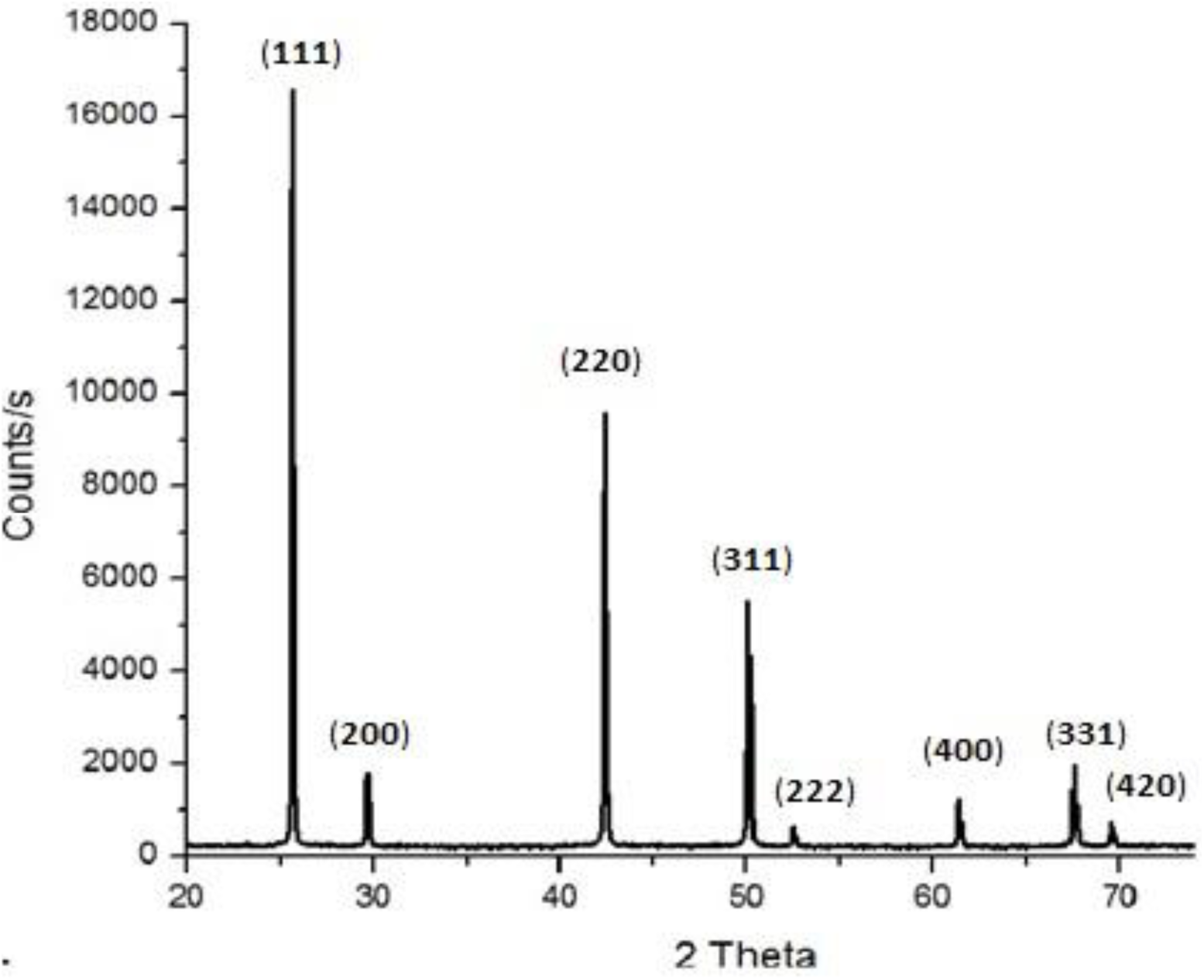
XRD OF CuI NANOPARTICLE.

**FIGURE 13.**
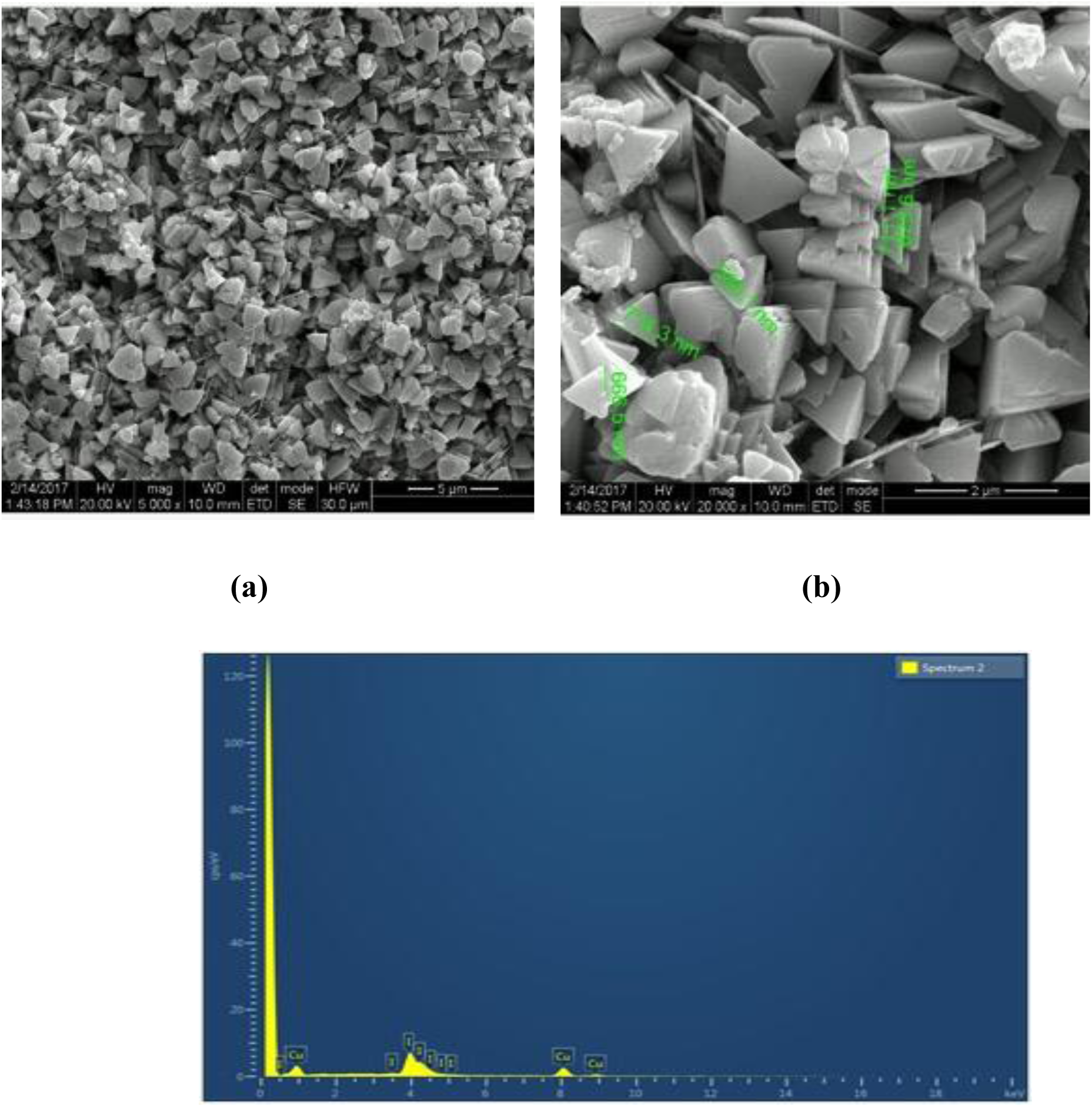
SEM (a) 5,000 X and (b) 20,000 X magnifications AND EDAX OF CuI NANOPARTICLE (Archana *et al.*, 2019)

### DEGRADATION OF AZO DYES BY CuI NANOPARTICLE

This is the first study of CuI nanoparticle in azo dye degradation before which 95% of cadmium adsorption was reported using Copper Iodide nanoparticle. The present study on degradation of azo dyes using CuI nanoparticle resulted in 15.835% for mixed dye, 16.9% for RBL, 3.262 % for RR and 16.745 % for RB of degradation of azo dyes at the end of 1 hour. The absorbance increased, which can be due to the limited ability of the CuI to adsorb dyes. The use of Strontium titanate in direct and reactive dyes resulted in increased degradation with increase in the Strontium titanate concentration (Karimi *et al.*, 2014).

### COMBINED METHOD FOR AZO DYES DEGRADATION

The degradation of azo dyes by the laccase enzyme, photocatalysis and using CuI nanoparticle are not sufficient for degrading the higher concentration of azo dyes. An effort to enhance the process of azo dye degradation was attempted by immobilizing laccase enzyme in CuI nanoparticle as shown in figure 14. The EDAX shows the presence of Cu, I, C, O. The presence of K is due to usage of potassium phosphate buffer during the immobilization process. The degradation of RR, RB, RBL and mixed azo dyes was increased by immobilizing Laccase enzyme in CuI nanoparticle. This method enhanced the percentage of degradation to 29.1%, 75.23%, 75% and 62.35% of RR, RB, RBL and Mixed azo dye of 100 ppm at 60^th^ minute. The HPLC analysis of the RR shows the presence of single peak at RT 2.4. Figure 15 (a) and the figure 15(b) shows the degraded RR of less intense peak at 3.4 RT. The RB dye showed an intense peak at RT 2.2, a small peak at RT 3 for non-degraded RB (Figure 16 a) shifted to RT 4.2 and the intense peak was significantly reduced at RT 2.5 after degradation by combined method as shown in figure 16 (b). The RBL dye (control) showed peaks at 2.5 and 6.1 RT (Figure 17 (a)) and the degraded RBL dye formed a peak at RT 3.4 (Figure 17 b). The mixed dye showed an intense peak as shown in figure 18 (a) at RT 2.3 and 3.0. The peaks after the degradation of mixed azo dye was similar to the RBL dye which shows that the RBL dye in the mixed azo dye was degraded efficiently than RB and RR dye (Figure 18 b). The GC-MS analysis of mixed azo dye by combined method resulted in the presence of 2-propen-1-amine-N-2propenyl, Benzoic acid, Naphthalene, 3-Butanynl benzene, 2-(m-tolyloxy) propionic acid, 4-chloro-N-(2-pyridinyl methylene) Benenanine, 9-Acetyl phenanthrene, Azacylcohexan-2-carboxylic acid amide 1-boc-N-methyl, Quinone 2,4-dinitrophenylhydrozone and Catechin (Figure 19). The results obtained are similar to Maulin Shah., (2013) on the degradation of Reactive Black dye by combining the process of photocatalytic and biological method showing 72% RB of 100 ppm. Biological degradation showed 85% of degradation of RB while in combined process the degradation percentage was found to be 74% without aromatic amine formation. The combined process enhanced the degradation of individual azo dyes compared to mixed azo dyes. The percentage of degradation is also higher compared to the biological and physical method. Thus, the metabolites produced by Biological and combined method of degradation of reactive azo dyes are different, therefore presumed to have different pathways.

**FIGURE 14.**
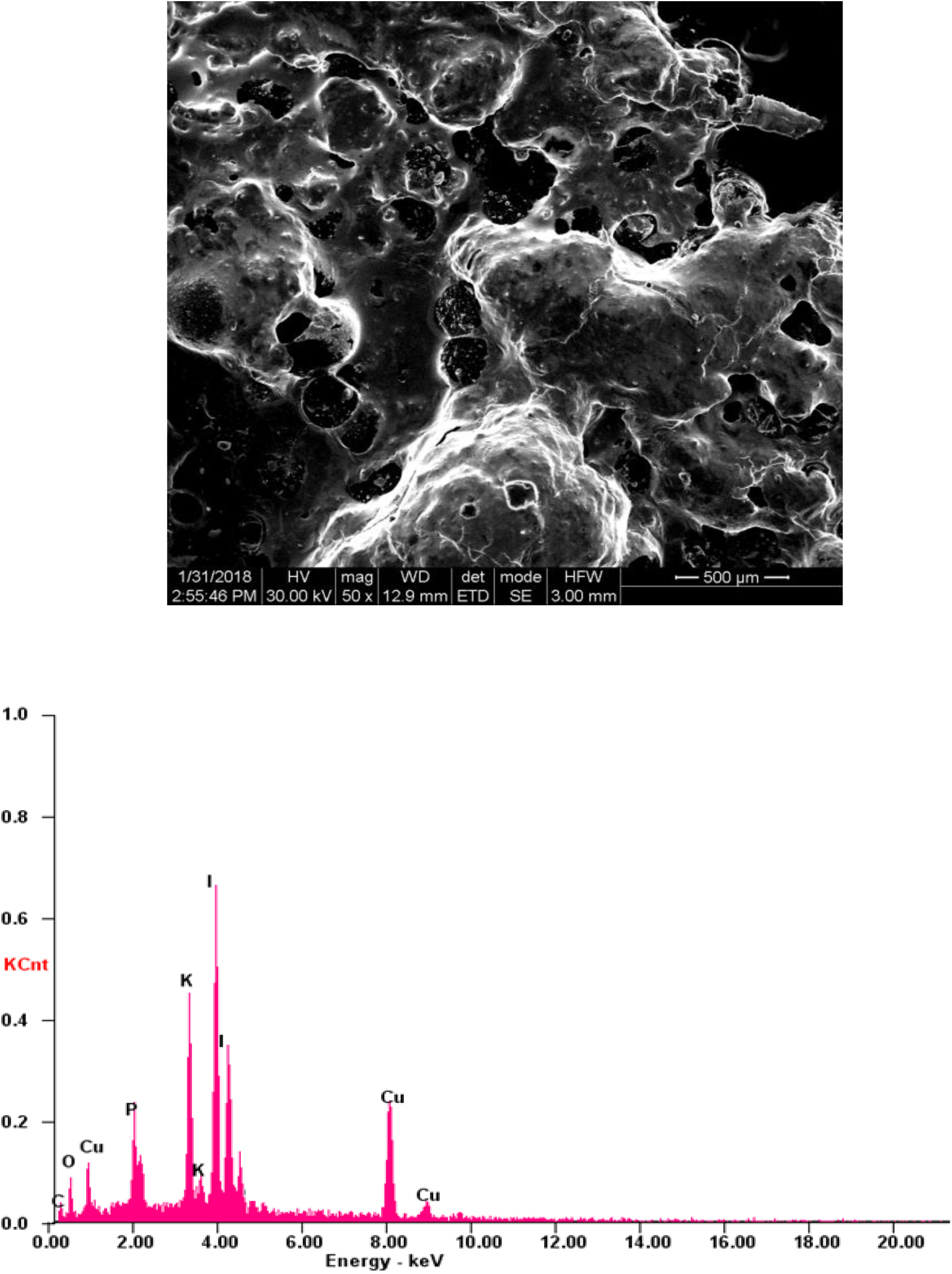
SEM AND EDAX OF LACCASE ENTRAPPED CuI NANOPARTICLE.

**FIGURE 15.**
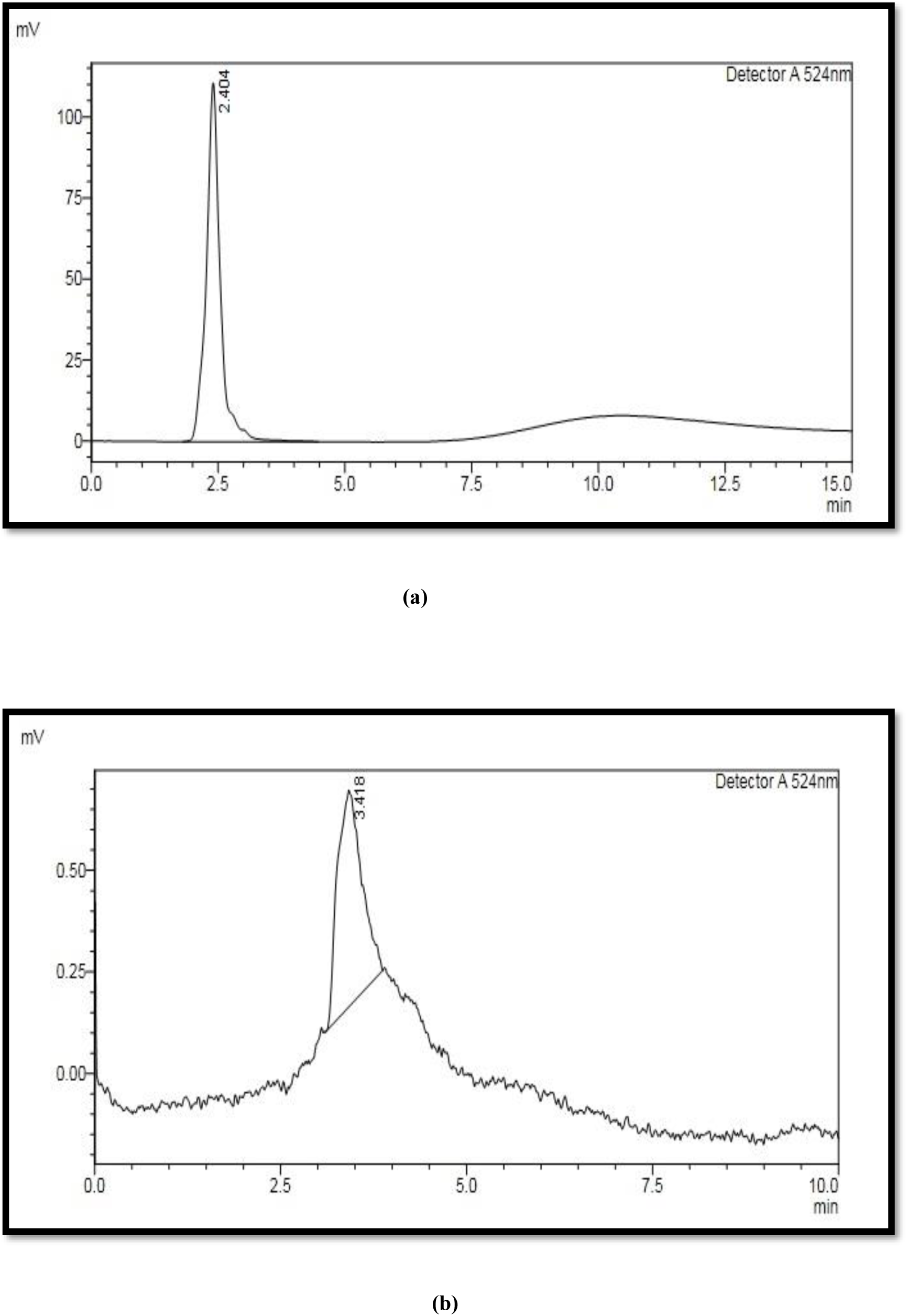
HPLC CHROMATOGRAM OF RR (a) CONTROL (b) DEGRADED BY COMBINED METHOD.

**FIGURE 16.**
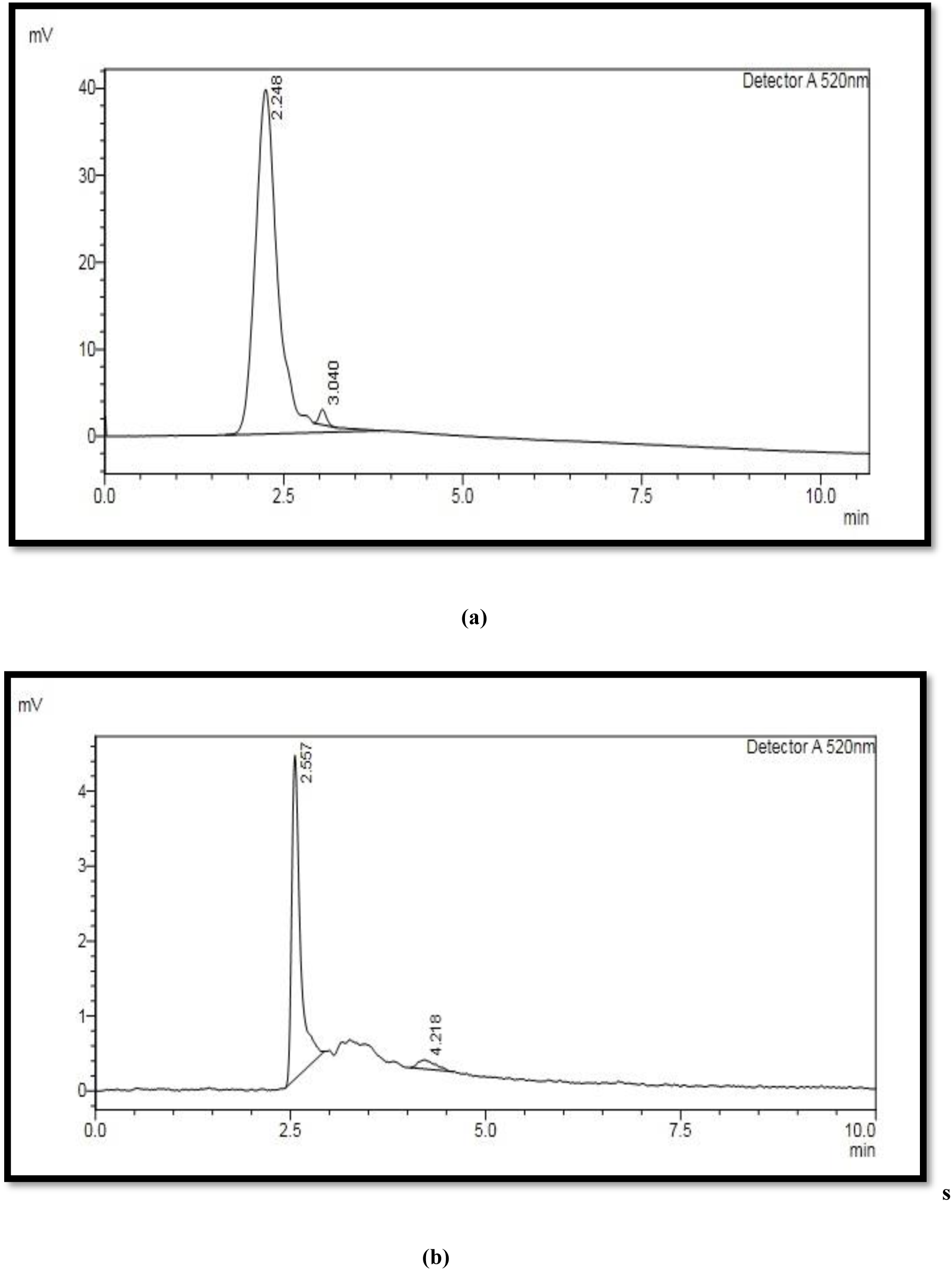
HPLC CHROMATOGRAM OF RB (a) CONTROL (b) DEGRADED BY COMBINED METHOD.

**FIGURE 17.**
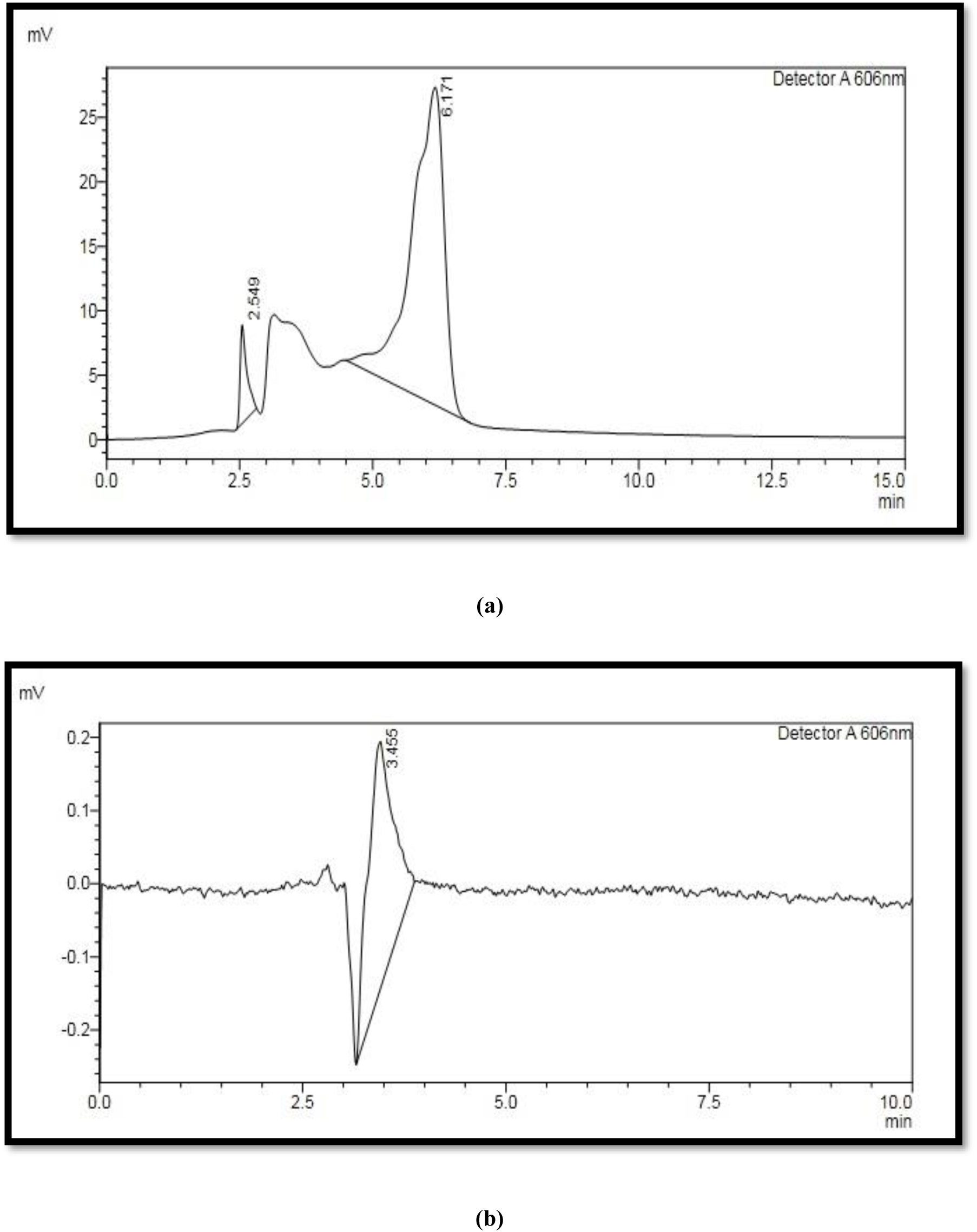
HPLC CHROMATOGRAM OF RBL (a) CONTROL (b) DEGRADED BY COMBINED METHOD.

**FIGURE 18.**
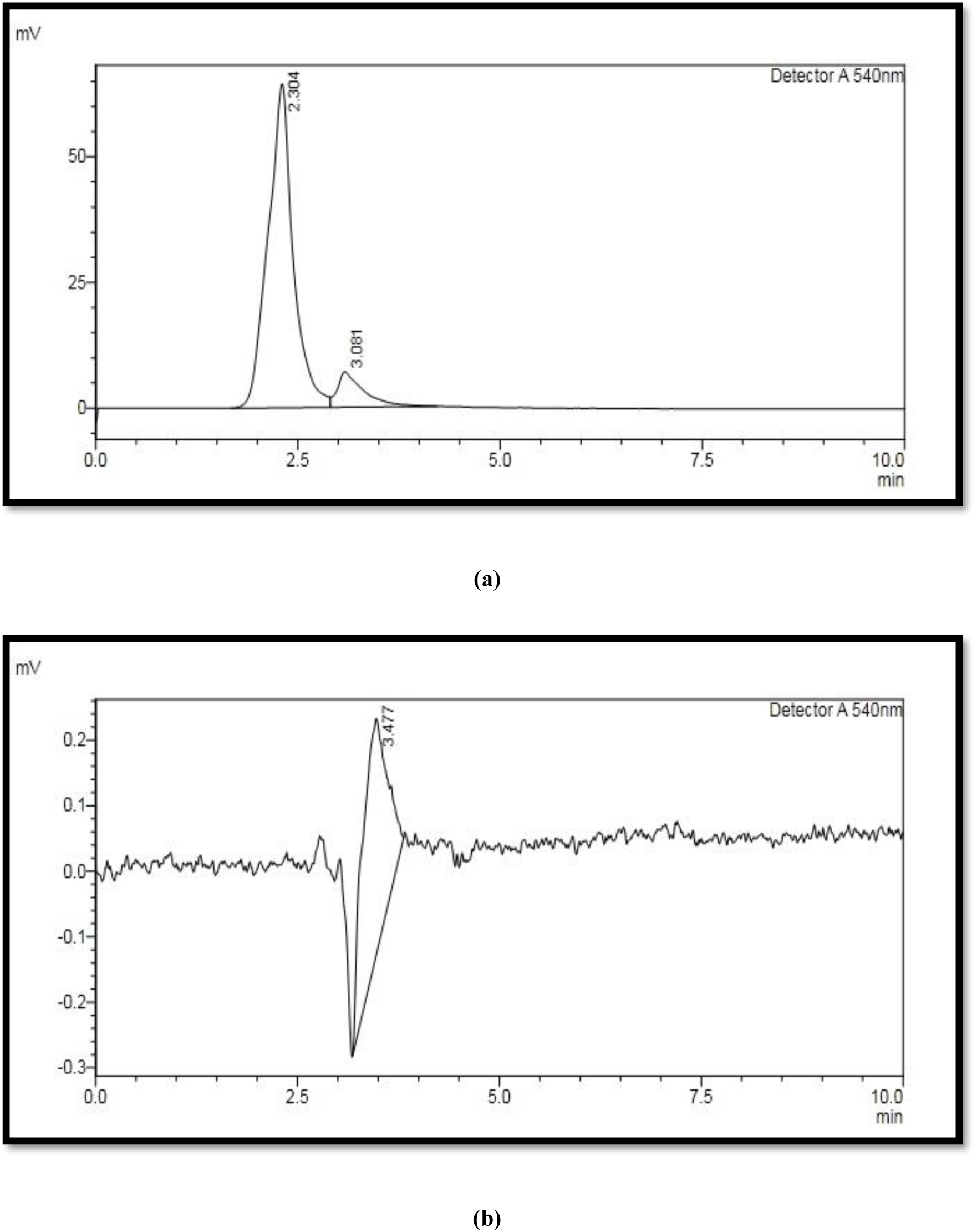
HPLC CHROMATOGRAM OF MIXED AZO DYE (a) CONTROL (b) DEGRADED BY COMBINED METHOD.

**FIGURE 19.**
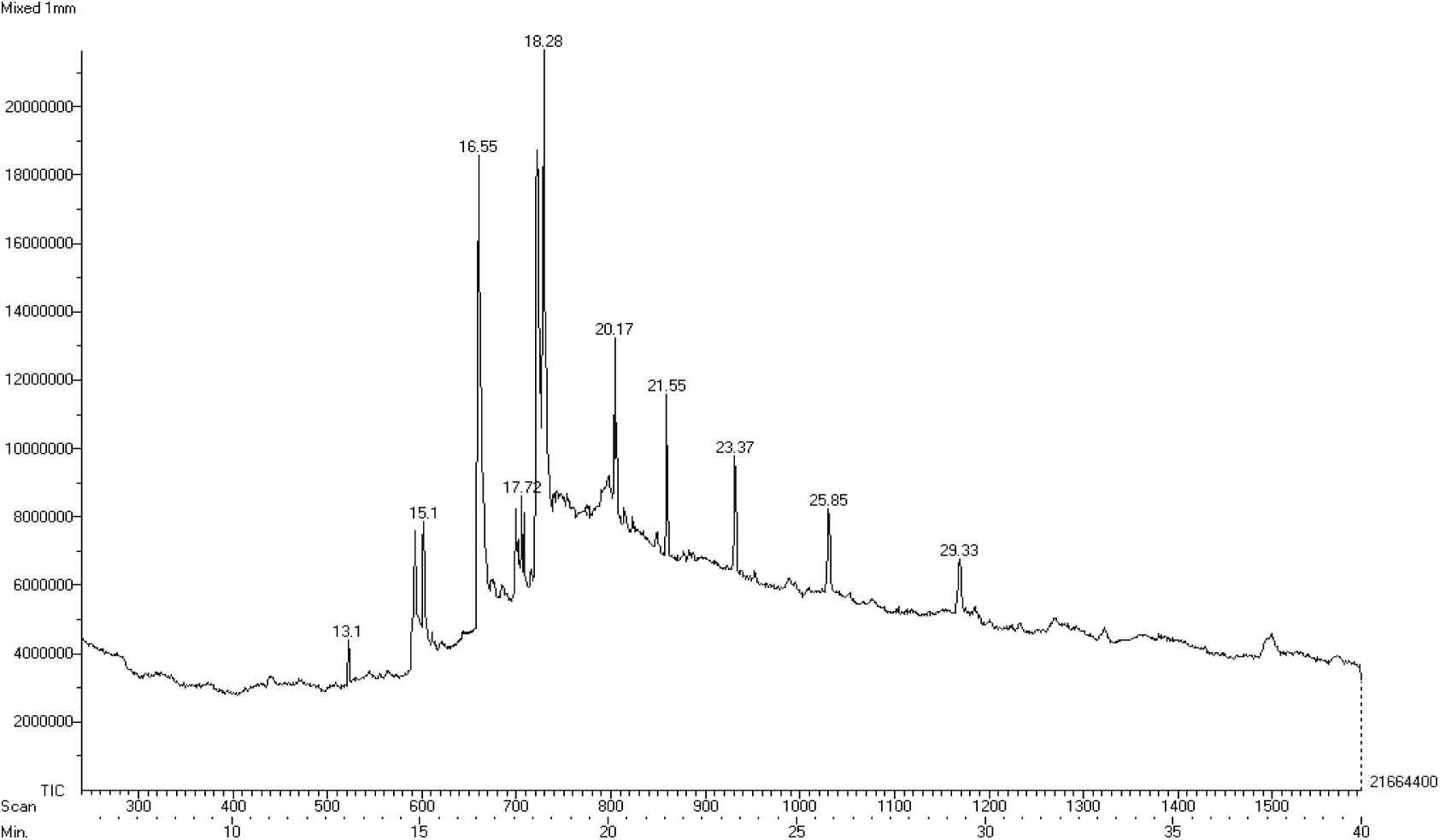
GC-MS SPECTRUM OF MIXED AZO DYE DEGRADED BY COMBINED METHOD.

### PHYLOGENETIC ANALYSIS OF *Pseudomonas mendocina*

The 16s RNA sequences of DY2KVG was blasted and it was found to be *Pseudomonas mendocina* (Genbank Accession number MK346304). The figure 20 shows the evolutionary history which was inferred using the UPGMA method (Sneath and Sokal 1973). The optimal tree with the sum of branch length = 1759.30814508 is shown. (next to the branches). The evolutionary distances were computed using the Maximum Composite Likelihood method (Tamura *et al.*, 2004) and are in the units of the number of base substitutions per site. The analysis involved 11 nucleotide sequences. Codon positions included were 1st+2nd+3rd+Noncoding. All positions containing gaps and missing data were eliminated. There were a total of 944 positions in the final dataset. Evolutionary analysis were conducted in MEGA7 (Kumar *et al.*, 2016).

**FIGURE 20.**
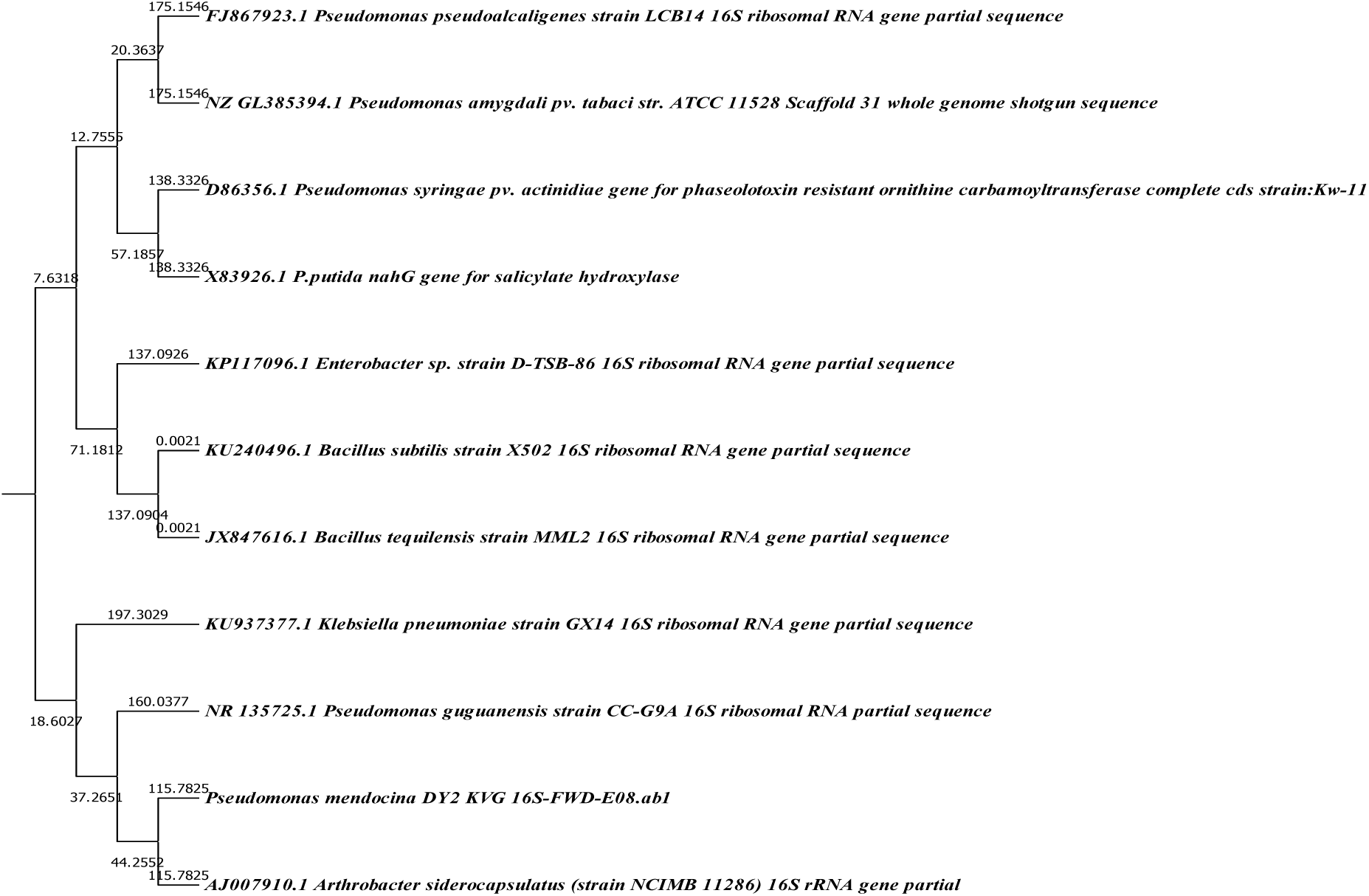
PHYLOGENETIC ANALYSIS OF *P.mendocina*.

#### *In silico* Analysis

From the Ramachandran plot evaluation using Rampage server, the protein model was found to have 92.7% of amino acids in the favoured region. Following amino acids were predicted to be the active site amino acids using CastP server: GLN63, PRO64, ALA65, TRP66, LEU67, VAL74, VAL75, ALA76, GLU77, ALA78, ASP79, PRO80, ALA81, THR82, VAL83, ILE84, ASN89, SER95, ILE96, and ALA97. From the molecular docking analysis, the average binding energies between the laccase and Reactive Black, Reactive Red and Reactive Brown were found to be -9.17 kcal/mol, -7.19 kcal/mol, and -8.57 kcal/mol respectively. The amino acids involved in the binding and degradation of dyes are shown in Figure 21 a, b, and c. The binding between the protein and the compound are stronger with lesser AutoDock binding energy (Morris *et al.*, 2009). From the docking results, we observed that the strongest and weakest binding of laccase was with Reactive Black and Reactive Red respectively. The binding between laccase and Reactive Brown was found to be in between the Reactive Red and Black. The strongest binding observed between the laccase, and Reactive Black could be the reason for the higher degradation of Reactive Black that was seen in our experimental findings.

**FGURE 21.**
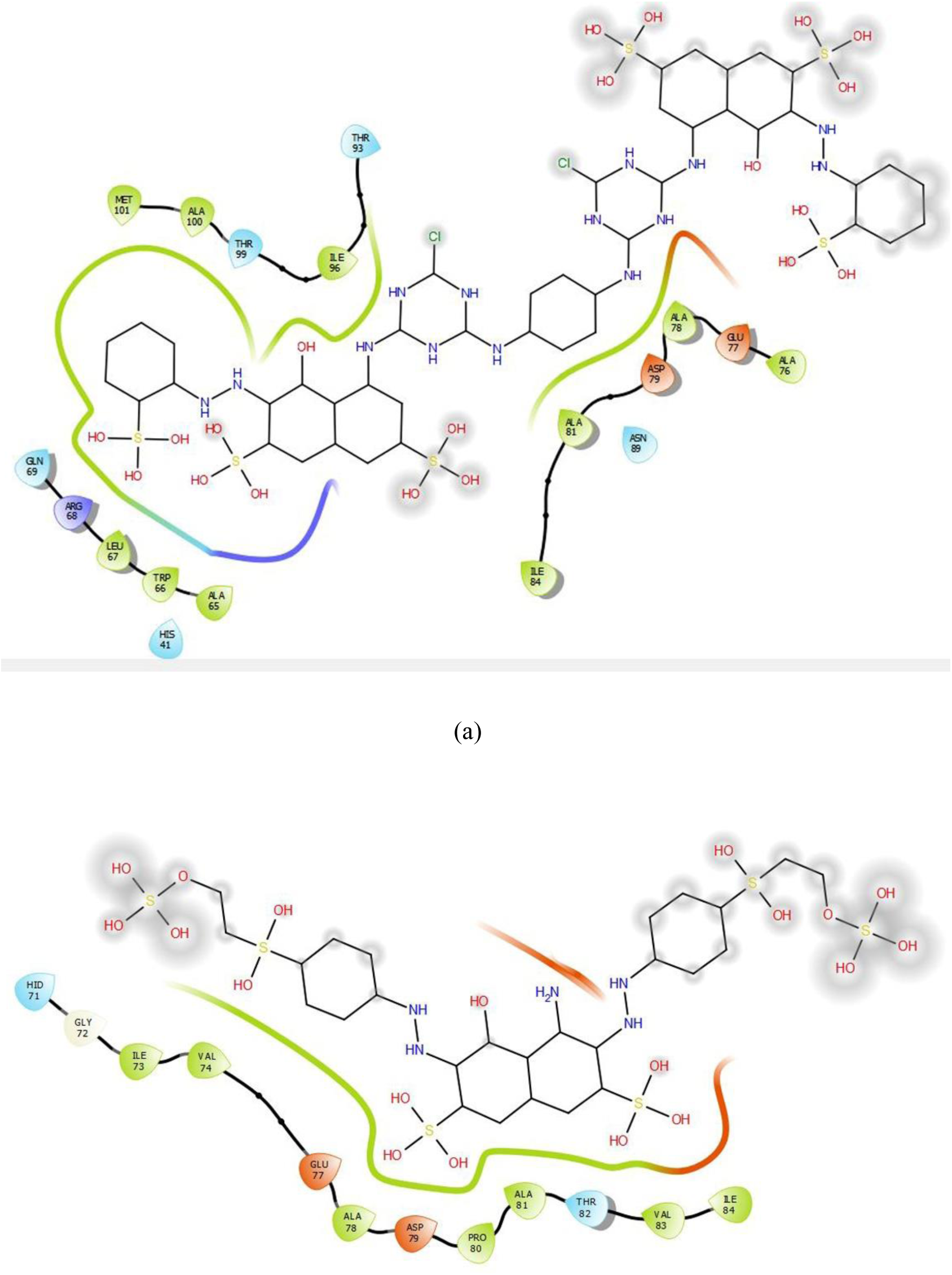
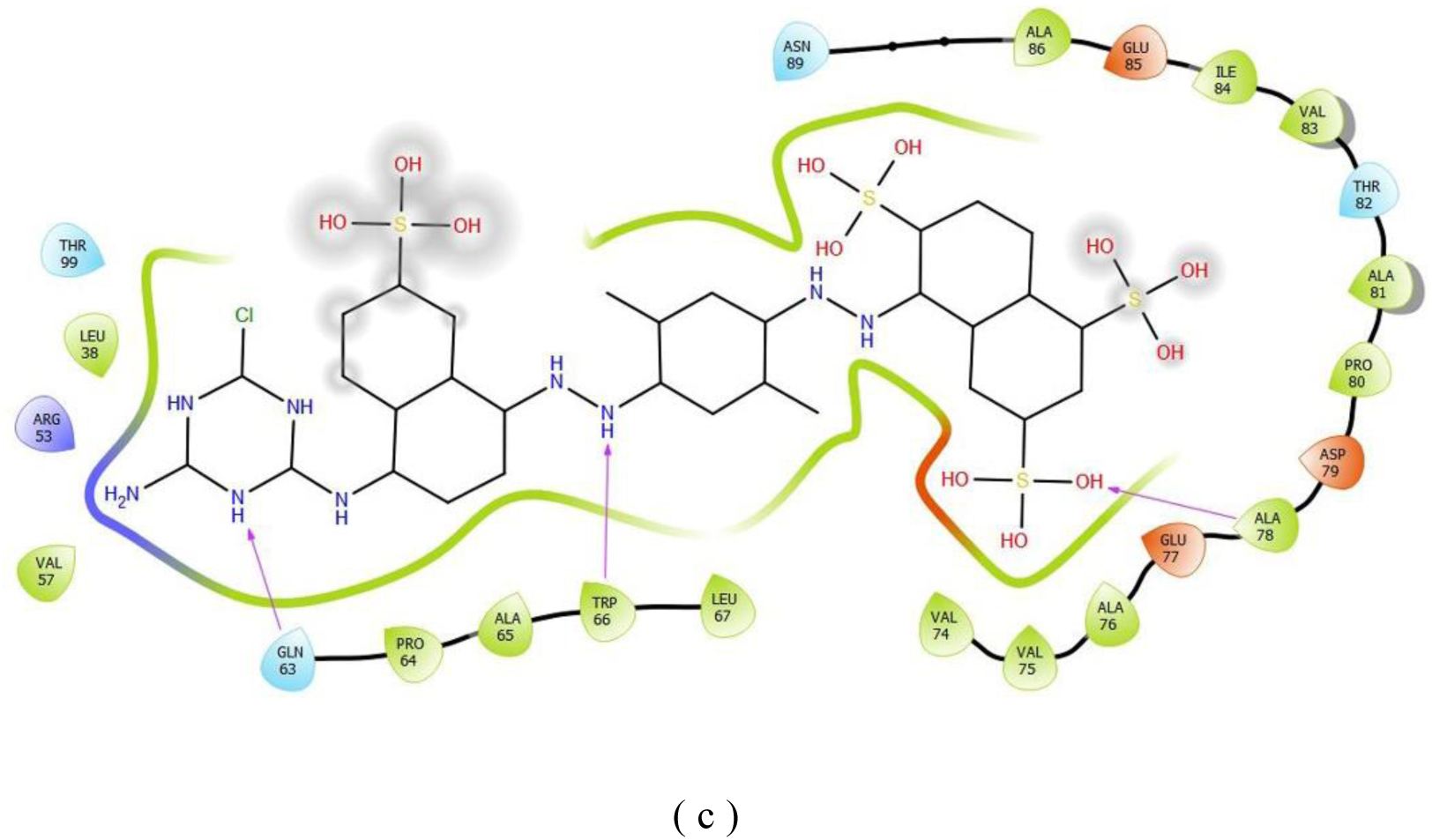
AMINO ACIDS BINDING AND DEGRADATION OF. **(a)RR DYE (b) RBL DYE (c) RB DYE**

## CONCLUSION

The degradation of azo dyes by laccase enzyme produced by *Pseudomonas mendocina,* an alkaliphilic bacterial strain resulted in 58.46% of degradation of mixed azo dye. The photocatalytic method was found to be less effective compared to biological method as it resulted in 15.98% of degradation of mixed azo dye at the end of 60^th^ minute. The CuI nanoparticle synthesised from *Hibiscus rosa sinensus* flower resulted in 15.835% degradation of mixed azo dye. Thus, the degradation of mixed azo dye was enhanced by immobilizing laccase enzyme in copper iodide nanoparticle which resulted in 62.35% of mixed azo degradation at the end of 60^th^ minute. The pathway of degradation of azo dyes using laccase enzyme, photocatalysis, CuI nanoparticle and combined method are different as the percentage of azo dyes (RR, RB, RBL and Mixed dyes) varies. In combined method the percentage of mixed dye degradation is in par with RB and RBL. While the percentage of degradation of individual azo dyes (RR, RB and RBL) are very less compared to the degradation of mixed azo dye by laccase, photocatalysis and CuI nanoparticle. The reusability of the free laccase enzyme and the combined CuI and laccase enzyme are yet to be studied. The kinetics of the degradation of individual dyes are in study to determine the order of reaction involved in the azo dye degradation by the enzymatic and combined method.

## ACKNOWLEDGEMENT

We would like to thank UGC-MRP for funding this project. We would like to thank CRIST LAB, Stella Maris College. We would like to extend our gratitude to SAIF IITM for SEM analysis.

## REFERENCES

1. Adosinda M Martins, Nelson Lima, Armando J D Silvestre and Maria Jao (2003) Comparative study of fungal degradation of single or mixed bioaccessible reactive azo dyes. Chemosphere 967–973

2. Alain K, Joël Q, Françoise L, Patricia P, Philippe C, Gérard R, Valérie C Marie-Anne CB (2002). Caminibacter hydrogeniphilus gen. nov., sp. nov., a novel thermophilic, hydrogen-oxidizing bacterium isolated from an East Pacific Rise hydrothermal vent. Intern. J. Systemat. Evolut. Microbiol. 52(4):1317–1323.

3. Alkesh I Shah and Jaydip B Jobanputra (2017) Production and Partial Purification of Laccase Produced by Bacillus species Isolated from Contaminated Soil. International Journal of Innovative Research in Science, Engineering and Technology 17862–17869

4. Andrea Zille, Barbara Gornacka, Astrid rehorek, Artur Cavaco Paulo (2005) Degradation of azo dyes by *Tremates villosa* laccase over long periods of oxidative conditions. Applied Environment Microbiology 6711–6718.

5. Anushya Vijayakumar, Revathy Rajagopal (2016) Green Synthesis and Characterisation of Copper (I) Iodide nanoparticles using kidney bean seed extract and its anti-bacterial activity. International Journal of Scientific & Engineering Research

6. Archana K M, Yogalakshmi D and Revathy Rajagopal (2019) Application of green synthesized nanocrystalline CuI in the removal of aqueous Mn(VII) and Cr(VI) ions. SN Applied Sciences.

7. Arka Mukhopadhyay, Anjan Kumar Dasgupta, Krishanu Chakrabarti (2013) Thermostability, pH stability and dye degrading activity of a bacterial laccase are enhanced in the presence of Cu2O nanoparticles. Bioresource Technology 127 25–36.

8. Biasini M, Bienert S, Waterhouse A, Arnold K, Studer G, Schmidt T, … Schwede T (2014) SWISS-MODEL: modelling protein tertiary and quaternary structure using evolutionary information. Nucleic Acids Research, 42 (Web Server issue), W252–8.

9. Binkowski, T. A., Naghibzadeh, S., & Liang, J. (2003) CASTp: Computed Atlas of Surface Topography of proteins. Nucleic Acids Research, 31(13) 3352–3355

10. Bordenstein, S. 2008. Microbial life in alkaline environments. http://serc.carleton.edu/microbelife/extreme/alkaline/index.html.

11. Dwakar V, Jadhav U and Govindwar S P (2008) Biodegradation of disperse textile dye brown 3REL by newly isolated *Bacillus Sps* VUS. Journal of applied microbiology 14–24

12. Fabbrini M, Glli, C and Gentili P (2002) Comparing the catalytic efficiency of some mediators of Laccase. J. Mol.Catal.B., Enzym 16 213–240

13. Harshad Lade, Sanjay Govinwar and Diby Paul (2015) Mineralization and detoxification of the carcinogenic azo dye congo red and real textile effluent by a polyurethane foam immobilized microbial consortium in an upflow column bioreactor. Int.J.Environ.Res.Public Health 12(6): 6894–6918.

14. Indubala E, Dhanasekar M, Sudha V, Revathy R, Venkataprasad Bhat S, Harinipriya S et al (2018) L-Alanine capping of ZnO nanorods: increased carrier concentration in ZnO/CuI heterojunction diode. RSC Adv 8:5350–5361. https://doi.org/10.1039/C7RA12385J

15. Jiang et al., (2011) Cauliflower-like CuI nanostructures: Green synthesis and applications as catalyst and adsorbent, Materials Science and Engineering B 176 1021–1027

16. Joshni T, Chacko Kalidass (2011) Subramaniam. Enzymatic degradation of azo dyes-a-review. Int.Jounal.of.envo.sci 0976–4402

17. Kalme, S., Parshetti, G. K., Jadhav, S. U and Govindwar, S. P. 2007. Biodegradation of Benzidine Based Dye Direct Blue-6 by *Pseudomonas desmolyticum NCIM 2112*. Bioresource Technology, 98: 1405.

18. Kobayashi H, Rittmann B E (1982) microbial removal of hazardous organic compounds, Environ. sci. technol. 16 170–183.

19. Kumar Raven GN and Sumangala K Bhat (2012) Fungal degradation of azo dye-Red 3BN and optimization of physic-chemical parameters. International journal of environmental sciences 17–24.

20. Lavanya C, Rajesh Dhankar, Sunil Chikars and Sarita Sheoran – Degradation of toxic dyes – a review, Int.j.curr.microbiol.app.sci. 2014.3(6):189–199

21. Loghman Karimi, Salar Zohoori, Mohammad Esmail Yazdanshenas (2014) Photocatalytic degradation of azo dyes in aqueous solutions under UV irradiation using nano-strontium titanate as the nanophotocatalyst. Journal of Saudi Chemical Society 18, 581–588

22. Lovell, S. C., Davis, I. W., Arendall, W. B., de Bakker, P. I. W., Word, J. M., Prisant, M. G., … Richardson, D. C. (2003). Structure validation by Calpha geometry: phi,psi and Cbeta deviation. Proteins 50(3) 437–450 https://doi.org/10.1002/prot.10286

23. Luciana Pereira, Ana V Coelho, Cristina A Viegas, Margarida M Correia dos Santos, Maria Paula Robalo, and Lígia O Martins (2009) Enzymatic biotransformation od azo dye suan orange G with bacterial cotA – laccase, Journal of biotechnology 139 68–77.

24. Luis A Ramírez-Montoya, Virginia Hernández-Montoya, Miguel A Montes-Morán, Juan Jáuregui-Rincón and Francisco J Cervantes (2015) Decolourisation of dyes with different molecular properties using free and immobilized laccases from *Trametes versicolor*. Journal of molecular liquids 30–37.

25. Madhuri M Sahasrabudhe, Rijuta G Saratale Ganesh D Saratale and Girish R Pathade (2014) Decolourisation and detoxification of sulphonated toxic diazo dye C.I direct red 81 by Enterococcus faecalis YZ 66. J. of envi health sci and engineering 12:151.

26. Maulin P Shah (2013) Combined Application of Biological-Photocatalytic Process in Degradation of Reactive Black Dye: An Excellent Outcome. American Journal of Microbiological Research 92–97.

27. Mohamed Neifar, Habib Chouchane, Mouna Mahjoubi, Atef Jaouani, Ameur Cherif (2016) Pseudomonas extremorientalis BU118: a new salt-tolerant laccase-secreting bacterium with biotechnological potential in textile azo dye decolourization. Biotech 6:107

28. Morris, G. M., Huey, R., Lindstrom, W., Sanner, M. F., Belew, R. K., Goodsell, D. S., & Olson, A. J. (2009) AutoDock4 and AutoDockTools4: Automated docking with selective receptor flexibility. Journal of Computational Chemistry 30(16) 2785–2791.

29. Morris, G. M., Huey, R., Lindstrom, W., Sanner, M. F., Belew, R. K., Goodsell, D. S., & Olson, A. J. (2009) AutoDock4 and AutoDockTools4: Automated docking with selective receptor flexibility. Journal of Computational Chemistry 30(16) 2785–2791.

30. N. Divya • A. Bansal • A. K. Jana (2013) Photocatalytic degradation of azo dye Orange II in aqueous solutions using copper-impregnated titania. Int. J. Environ. Sci. Technol. 10:1265–1274

31. Rakesh K Soni, P. B. Archarya and H.A. Modi (2015) Elucidation of biodegradation mechanism of Recative Red 35 by Pseudomonas aeruginosa ARSKS20. IOSR-JESTFT 31–40.

32. Roberto Comparelli, Elisabetta Fanizza, M. L. Curri and Angela Agostiano (2005) UV induced photocatalytic degradation of azo dyes by organic capped ZnO nanocrystals immobilized onto substrates. Applied catalysis B: Environmental 60 1–11.

33. Salleh M A M, Mahmoud M K, Karim W A, Idiris A (2011) Cationic and anionic dye absorption by agricultural solid wastes: a comprehensive review, Desalination 1–13.

34. Saratale RG, GD Saratale, DC Kalyani, IS Chang and SP Govindwar (2011) Bacterial decolourisation and degrdataion of azo dyes – A Review. Journal of Taiwan institute of Chemical Engineers 42:138–157.

35. Saratale RG, GD Saratale, DC Kalyani, IS Chang and SP Govindwar (2011) Bacterial decolourisation and degrdataion of azo dyes – A Review. Journal of Taiwan institute of Chemical Engineers 42:138–157.

36. Satadru Pramanik and Sujata Chaudhuri (2018) Laccase Activity and Azo Dye Decolorization Potential of *Podoscypha elegans*. MYCOBIOLOGY 46(1) 79–83

37. Shyamala Gowri R, Vijayaraghavan R and Meenambigai P (2014) Microbial degradation of azo dyes-review. Int. j.curr.microbiol.app.sci. 3(3):421–436.

38. Sujatha Boopalan, Veena Gayathri and Vidhya Vasudhevan (2017) Alkaliphilic degradation of mixed reactive dyes (C.I. RY 125 and C.I. RB 52) by a moderately alkaliphilic bacterial consortium – optimization and bench scale studies. Basic Research Journal of Microbiology 2354-4082 4(5) 47–63.

39. Tarun A, Rachana S (2012) Bioremedial potentials of a moderately halophilic soil bacterium. JPBMS 19:1–6.

40. Tavakoli F, Salavati-Niasari, M, Mohandes, F (2013) Green Synthesis of Flower-like CuI Microstructures Composed of Trigonal Nanostructures Using Pomegranate Juice. Mater. Lett 100 133–136

41. Vakili M, Rafatullah M, Salamatinia B, Abdullah A Z, Ibrahim M H, Tan K B, Gholam Z, Amouzgar P (2014) Application of chitosen and its derivatives as absorbants for dye removal from water and waste: a review. Carbohydr. Polym 113 115–130.

42. Yi Jiang, Shuyan Gao, Zhengdao Li, Xiaoxia Jia, Yanli Chen (2011) Cauliflower-like CuI nanostructures: Green synthesis and applications as catalyst and adsorbent. Materials Science and Engineering B 1021–1027.

43. Zahra Asadgol, Hamid Forootanfar, Shahla Rezaei and Mohammad Ali Faramarzi (2014) Removal of phenol and bisphenol-A catalysed by laccase in aqueous solution. Journal of Environmental Health science and engineering 12; 93.

